# Viral-structured models of dividing cells shows cell-virus coexistence via viral load partitioning between cell progeny

**DOI:** 10.1101/2020.11.17.384842

**Authors:** Hugh Z. Ford

## Abstract

The present study develops and analyses a system of partial differential equations that model a single population of dividing cells infected by lytic viruses in a closed system. This mean-field model stratifies cells by cell size (continuous) and number of virus particles per cell (discrete) to couple the cell cycle and the lytic cycle under mass conservation. We present numerical solutions to the mean-field model and an equivalent stochastic model for parameter values representative of *Escherichia Coli* and lytic bacteriophages such as Escherichia virus T4. This analysis suggests that dividing cells and lytic virus populations in isolation can coexist in the absence of evolutionary, ecological and biochemical processes. Coexistence emerges because viral load dilution via cell growth and viral load partitioning via cell division both counteract viral load growth via viral synthesis and hence cell death by lysis. Furthermore, we analytically determine the quasi-steady state solution of the mean-field model in the continuum limit with respect to viral loads. From this solution we derive a condition for cell-virus coexistence through viral load partitioning: that the product of the viral synthesis rate, cell lysis rate and the time between cell divisions must be less than the product of log(2) and the cell growth rate. Overall, the present study provides a theoretical argument for a stable relationship between cells and lytic viruses simply by virtue of cell growth and division.

## 1 Introduction

For over 1.5 billion years viruses have infected and coexisted with cells [1]. Once inside a cell, a virus typically replicates until the host dies by lysis, which then allows numerous viral progeny to infect new cells (see Figure 1A). This life cycle, called the lytic cycle, produces an unsustainable predator-prey relationship between viruses and their target cells. To avoid the collapse of both viral and cell populations, some viral species (e.g. *Escherichia virus Lambda*) can switch between the lytic cycle and a dormant and heritable state within the cell (lysogenic cycle) [2]. However, for lytic viruses, population collapse is avoided by the rapid evolution (due to large population sizes, selection pressures and mutation rates) of diverse biochemical and ecological strategies spread across large spatial domains [3–9]. Coevolutionary arm races emerge when mutant bacteria become dominant following the gain of a new mechanism that resists viral infection [10]; once dominant in the cell population, mutant viruses that bypass this new cellular mechanism subsequently become dominant in the virus population [11]. Through this evolutionary process, bacteria and viruses coevolve into phenotypically heterogeneous communities that form complex infection networks shaped by diverse ecological strategies such as predator-resistance, host-range trade-offs and bet-hedging [8, 12, 13].

**Figure 1:**
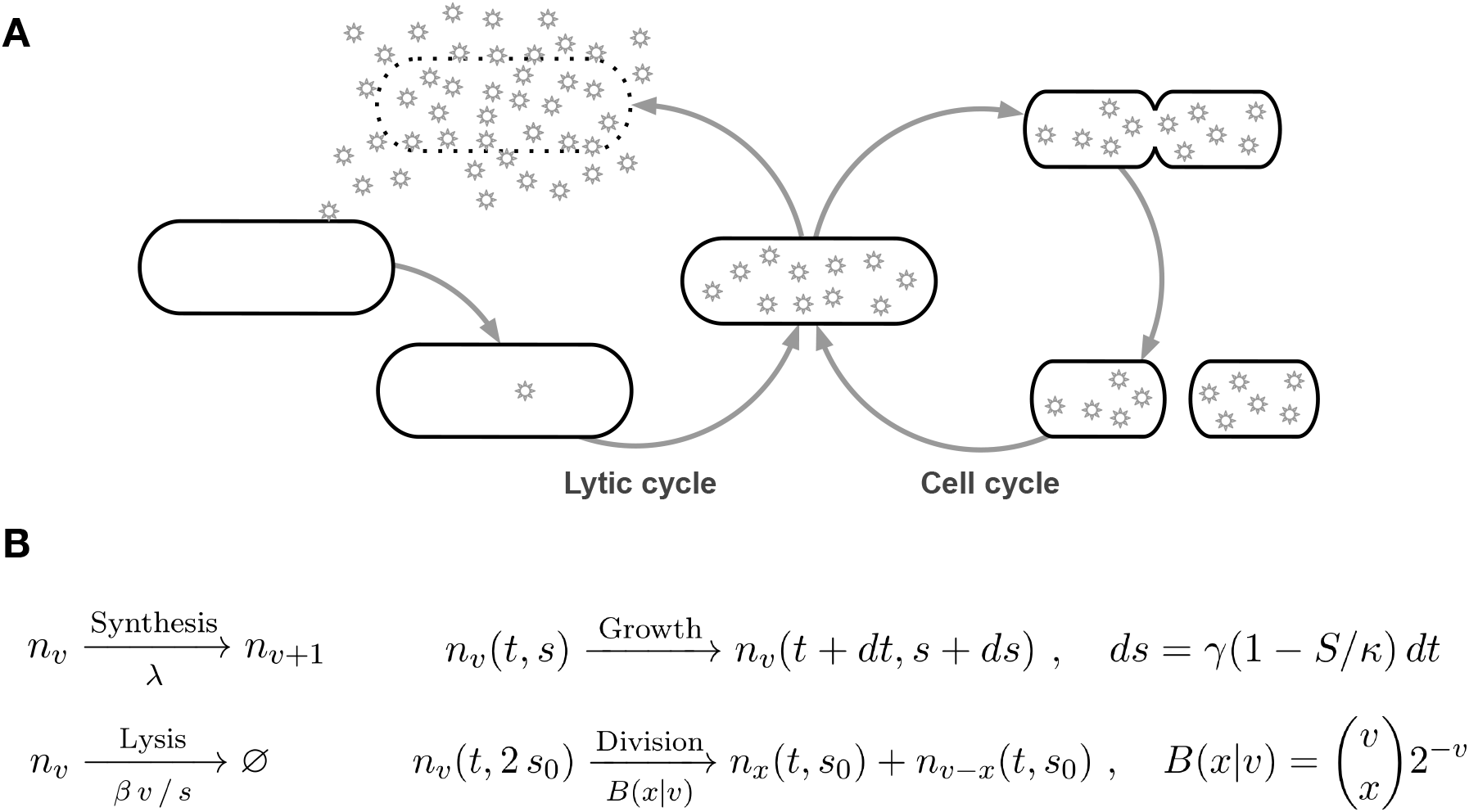
A schematic of passive coexistence between lytic viruses (small star) and bacterial cells (large ellipse). In the lytic cycle (left), cells are infected with an extracellular virus which replicates within the host cell until the cell dies by lysis. The lytic viruses that are released into the environment following cell lysis can repeat the cycle by infecting a new cell. In the cell cycle (right), cells grow in size and then divide to produce two daughter cells. During cell division the parent cell viral load is split between the two daughter cells. There are two fates of a virus-infected cell: cell death or division.

System of Lotka-Volterra equations (system of ordinary differential equation) resembling the SIR model (infectious disease transmission) are commonly used to mathematically model mean-field virus and cell population dynamics as a predator-prey system [1, 14–17]. Lotka-Volterra models compartmentalise cells and viruses into susceptible cell, infected cell and extracellular virus classes and predict cell-virus coexistence through the perpetual proliferation and infection of susceptible cells. These models successfully reproduce key features of infected cell population dynamics and are typically at the core of theoretical investigations on virus-cell coexistence [1, 18]. However, the use of Lotka-Volterra models often provides a narrow view of ecological systems. In this paper, we study cell and viral coexistence using a McKendrick–von Foerster model that couples cell and viral growth together.

We hypothesise that the coupling between the cell cycle and the lytic cycle enables coexistence between cells and viruses in the absence of evolutionary, ecological and biochemical processes (see Figure 1). Our reasoning is as follows. Cell division partitions the contents of a cell between cell progeny. As such, molecules that are synthesised at a rate lower than the cell growth rate dilute within proliferating cell populations [19]. It is thus conceivable that lytic viruses also dilute within proliferating cell populations such that proliferating cell populations continuously partition proliferating virus populations into viral loads that are smaller than the burst size. In this way, both the lysogenic cycle and the cell cycle promotes cell-virus coexistence via hereditary transduction of lysogenic and lytic viruses respectively.

The present study uses proof-of-concept mathematical modelling to test the verbal model above. We build upon existing partial differential equations (PDE) that model bacteria growth via the use of a structural variable *s* to track cell size [20] by incorporating another structural variable *v* that tracks the number of virus particle per cell. This approach produces an infinite system of first order hyperbolic partial differential equations (PDE) with non-local and periodic boundary conditions which accounts for cell-to-cell variance with respect to cell size and viral load [21] (as opposed to Lotka-Volterra/SIR models that consider the mean-field population dynamics of homogeneous/compartmentalised infected and uninfected cells). This system of equations models the time evolution of the distribution of cell size *s* and viral loads *v* among cells as cells grow in size and divide, and synthesise new virus particles and die by lysis. Viral loads concentrate due viral synthesis, dilute as cells grow and are partitioned between cell progeny during cell division. We assume that the cell lysis rate is proportional to the the number of virions per cell volume *v*/*s*, and that cells logistically grow in size and divide when cells increase in size by the minimum cell size [22]. We use this new model to show that perpetual viral synthesis and cell proliferation can stably maintain viral loads below excessive values that rapidly trigger cell death by lysis. As such, the cells decision to divide or die (and hence whether a population grows or decays in size) is determined by the relative rates of viral synthesis and cell growth. Our model suggests that cells and lytic viruses coexist when the mean division time (the time taken for a newborn cell to divide) is less than half the mean lysis time (the time between initial viral synthesis and cell death).

The present study is organised as follows. In section 2 we introduce the mean-field PDE model of cell and viral growth. This introduction is accompanied by the derivation of the model from first principles in Appendix A, derivation of an equivalent stochastic model in Appendix B, and derivation of model parameter values based on *Escherichia coli* and bacteriophages such as bacteriophage *T4* and λ. In section 3 we present and describe numerical time-dependent solutions of the mean field model and numerical simulations of the stochastic model. These results demonstrate cell-virus coexistence for typical parameter values associated with *E. coli* and bacteriophages. In section 4 we produce and describe analytic expressions for the long time distribution of viruses and size among cells during the exponential and stationary phase of cell population growth using a continuum approximation of the mean field model (where the mean burst size is assumed to be very large). This analysis allows us to determine the conditions for cell-virus coexistence in terms of the relationship between parameters of cell and viral growth. The supplementary material presents an age- and viral load-structured model which is representative of Eukaryotic cells and assumes that cells divide with a logistic rate following a fixed time period for cell growth and that cell lysis occurs instantaneously at a critical viral load. Model development and analysis mirrors that which is presented in the main text (i.e. derivation of a mean-field and stochastic model, numerical time-dependent solution and analytical expressions from the continuum approximation of the mean-field model). In section 5 we discuss the significance of our findings.

## 2 Model derivation

We have derived and analysed equivalent mean-field (outlined in Appendix A) and stochastic (outlined in Appendix B) model to study the coexistence of viruses and cells in a well-mixed and closed system. These models consider the time evolution of a single infected cell population *n_v_* (*t, s*) and keep track of cell size 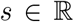 (*μ*m^3^) and viral load 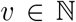 (number of virions) as cells: synthesise new viruses (which concentrates viral loads), grow logistically (which increases cell size and dilutes viral loads), die by lysis (which removes cells and releases viruses), and divide (which partitions the viral load and cell size of a single cell between two), as outlined in Figure 1A. The mass action kinetics of these processes are outlined in Figure 1B. Model derivation and parameterisation was motivated by bacteria and bacteriophages. We assume a constant viral synthesis rate and a cell lysis rate that is proportional to the viral concentration, *v*/*s* (number of virions *μ*m^−3^). Cell growth is modelled according to the ’adder’ mechanism of bacterial growth [22], which was expressed mathematically, and experimentally validated, in Ref. [20]. We assume that newborn cells have size *s* = *s*_0_ and dividing cells have size *s* = 2 *s*_0_. Coupling these processes together under mass conservation produces the following infinite system of viral load- and size-structured, first order hyperbolic partial differential equations (PDE) with non-local, periodic boundary conditions:

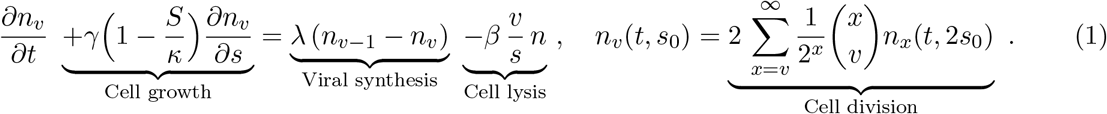

with *s*_0_ ≤ *s* ≤ 2 *s*_0_ and *v* = 0, 1,… (and *n*_−1_ = 0). We consider an initial condition where every cell is infected but does not contain any virions *n*_0_(0, *s*) = *σ*(*s*) (and *n_v_*(0, *s*) = 0, *v* > 0) where *σ*(*s*) is the initial cell size distribution. The total number of cells *N* = *N*(*t*), the collective volume of the cell population *S* = *S*(*t*) and the total number of intracellular virus particles *V* = *V*(*t*) are respectively defined as:

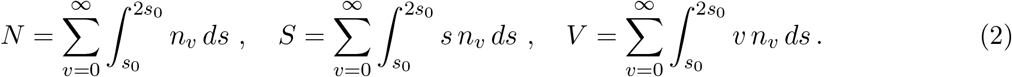

An alternative age- and viral load-structured model is derived and analysed in the supplementary material. This model is similar to equation (1) except that: (i) cell size is neglected and cell age is considered, (ii) cell division occurs at a logistic rate (dependent on the number of cells) following a delay phase following the time of parental cell division and (iii) cell death by lysis is assumed to occur instantaneously at the mean burst size.

## 3 Numerical results for bacteria-bacteriophage coexistence

The section presents and describes a numerical solution of the mean-field model (equation (1), Appendix A) and a numerical simulation of the stochastic model (Appendix B) using parameter values (shown in Table 1) that are representative of *E. coli* infected with bacteriophage T4 or λ (see Appendix C for derivation). The method of lines was used to numerically solve the mean-field model so as to preserve the jump discontinuity in the solution *n_v_* (*t, s*) across *s* (unlike discretisation methods such as Crank-Nicholson and upwind scheme). The Gillespie algorithm was used to simulate the stochastic model.

**Table 1:** Default parameter values for *E. coli* infected with bacteriophages such as T4 and λ, derived in Appendix B.

| Parameter | Symbol | Default value | Range | Dimension | Source |
| --- | --- | --- | --- | --- | --- |
| Maximum division rate | $\alpha$ | 2 | 0 - 3 | hr<sup>-1</sup> | [20] |
| Minimum division time | $\tau$ | 0.5 | 0.2 - 24 | hr | [20] |
| Newborn cell volume | $s_0$ | 1 | 0.3-3 | $\mu\text{m}^3$ | [20] |
| Cell growth rate | $\gamma$ | 2 | 1 - 10 | $\mu\text{m}^3 \text{ hr}^{-1}$ | $s_0/\tau$ |
| Virus synthesis rate | $\lambda$ | 200 | 10-1000 | particles hr<sup>-1</sup> | [1, 23] |
| Cell lysis rate | $\beta$ | 0.005 | 0.001 - 0.01 | particles | [24] |

### 3.1 Coexistence ancestral lineage

Figure 2A presents a single simulation of the stochastic model for small numbers of cells (carrying capacity *κ* = 30 *μ*m^3^) from an initial condition where there is one infected cell with minimum cell size *s* = *s*_0_ and no viral load *v* = 0. The cell and virus population are represented as a single ancestral lineage that tracks cellular: (i) viral load *v* (virions per cell), (ii) viral concentration *v*/*s* (virions per *μ*m^3^ per cell), (iii) size *s* (*μ*m^3^ per cell), (iv) age (via edge lengths), (v) division (branch points) and (v) death by lysis (terminal nodes). As such, this representation shows how infected cells can negate cell lysis through the dilution and partitioning of viral loads respectively via cell growth and cell division. Viral load dilution reduces the likelihood of cell lysis and hence increases the likelihood of cell replication (and viral load partitioning). Consequently, robust coexistence immediately follows infection when the extent of viral load dilution is greater than the extent of viral load concentration via virion synthesis. We note that the random partitioning (given by a binomial distribution) of viral loads between cell progeny produces cell-to-cell differences which, together with stochastic viral production and cell growth, can lead to different outcomes (i.e. death or division) for daughter cells [21] and causes the total number of cells to randomly fluctuate around a positive value.

**Figure 2:**
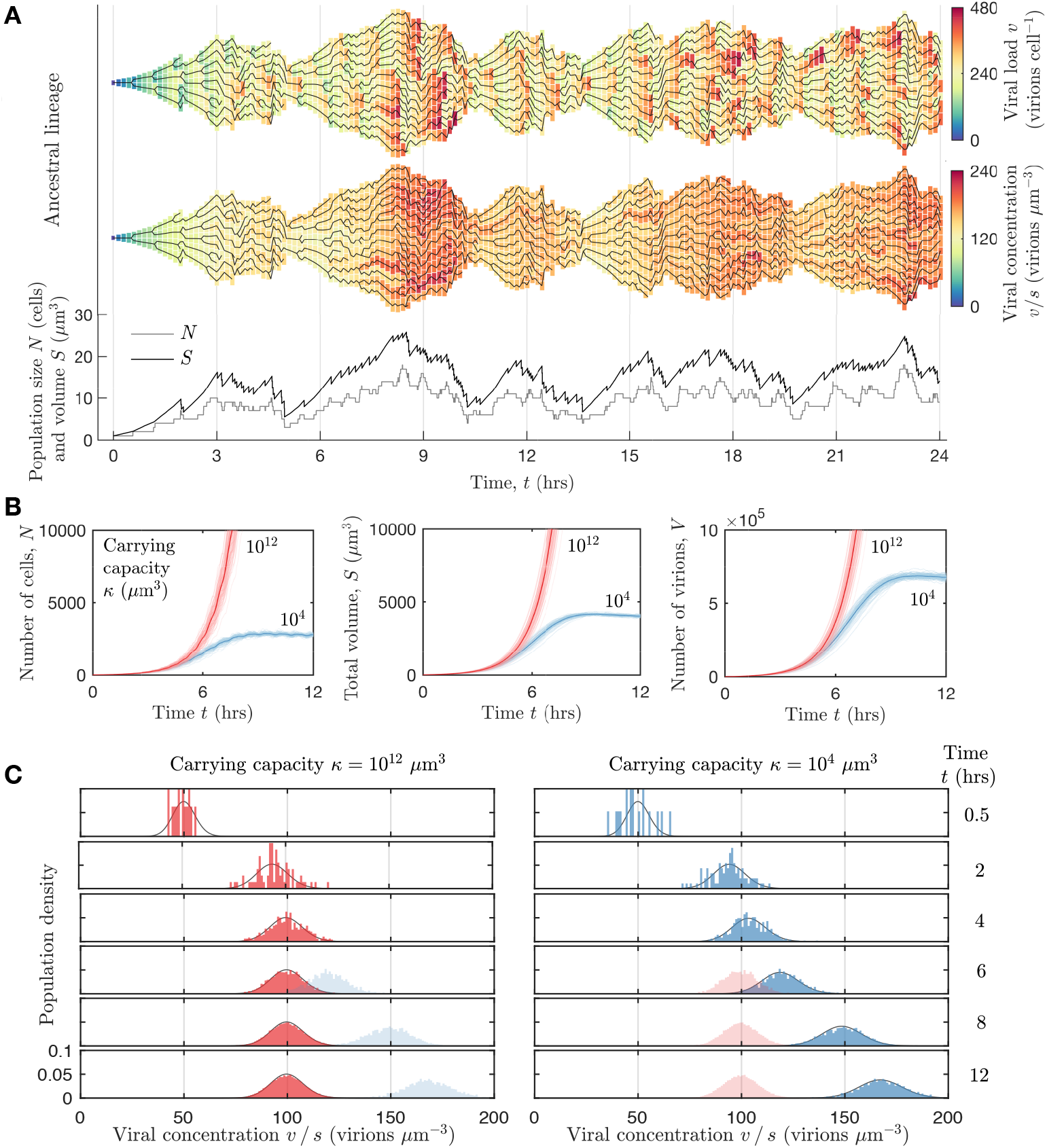
**A** A simulation of the stochastic model (Appendix A) with carrying capacity *κ* = 30 *μ*m^3^ and initially one infected cell with size *s* = 1 *μ*m^3^ and no viral load *v* = 0. Model parameter values are shown in Table 1 and include a cell growth rate variance of 1 *μ*m^3^ hr^−1^ and cell division size variance of 0.1 *μ*m^3^. Shown: the ancestral lineage (solid black line) tracking cell division and lysis overlaying the cell population represented as a set of cells (coloured blocks, stacked vertically) with size *s μ*m^3^ (block length) and viral load *v* (block colour, top lineage) and the viral concentration *v*/*s* (block colour, middle lineage) and the population size *N* (grey line) and volume *S* (black line) (bottom). **B** The mean total number of cells N (left), population volume *S* (middle) and total number of intracellular viruses (right) from 100 simulations of the stochastic model with carrying capacity *κ* = 10^4^ *μ*m^3^ (blue) and *κ* = 10^12^ *μ*m^3^ = 1 mL (red). **C** The distribution of viral concentration *v*/*s* among cells generated from (i) a single simulation of the stochastic model with carrying capacity *κ* = 10^4^ *μ*m^3^ (blue) and *κ* = 10^12^ *μ*m^3^ = 1 mL (red) and (ii) a numerical solution to the mean-field model (black line), see Figure 3.

### 3.2 Viral load- and cell size-distribution during exponential and stationary growth

Figure 2B presents the total number of cells *N*, total cell volume *S* and total number of intracellular virions *V* given by the mean of 100 simulations of the stochastic model for large numbers of cells (carrying capacity *κ* = 10^4^ *μ*m^3^ and *κ* = 10^12^ *μ*m^3^ = 1 mL). These simulations considered an initial condition where there are 10 cells whose cell size is randomly generated from values between *s* = *s*_0_ and *s* = 2 *s*_0_. Figure 2C presents the distribution of viral concentrations *v*/*s* (virions *μ*m^−3^) among large numbers of cells and contrasts the distribution profiles during the exponential phase and stationary phase of cell population growth (where the cell population is respectively much lower than, and close to, carrying capacity). During the exponential growth phase, constant cell growth counteracts constant viral synthesis such that viral concentrations are stably contained at values where cell lysis rates are less than cell division rates. Intercellular competition decelerates cell growth until the population reaches the stationary growth phase. A reduction in cell growth and viral dilution rates (while viral synthesis remains constant) concentrates viral loads and hence increases cell lysis rates. At the stationary phase of population growth the viral load are stably distributed among cells at a mean concentration that produces equal cell death and division rates.

### 3.3 Mean-field breather dynamics of cell-virus growth

Figure 3A shows the population size *N* and volume *S* during the exponential and stationary phase of cell population growth (which closely matches stochastic model output shown in Figure **??**). This solution has an initial condition where cell size is uniformly distributed across a cell population of size 1, *σ*(*s*) = 1/*s*_0_ (*s*_0_ ≤ *s* < 2*s*_0_ *μ*m^3^). The cell and virus population tends to a state where the total number of infected cells *N* oscillate over time [25]. As opposed to a predator-prey interactions [1], these oscillations arise in our model (equations (1) and (**??**)) because the distribution of viruses and size among cells *n_v_*(*t,s*) dynamically cycles between quasi-steady states (i.e. a breather solution [26]). Breather dynamics is underpinned by incremental increases in cell size *s* (cell growth) and viral load *v* (viral synthesis) and non-local halving (approximately) of both *s* and *v* due to cell division.

**Figure 3:**
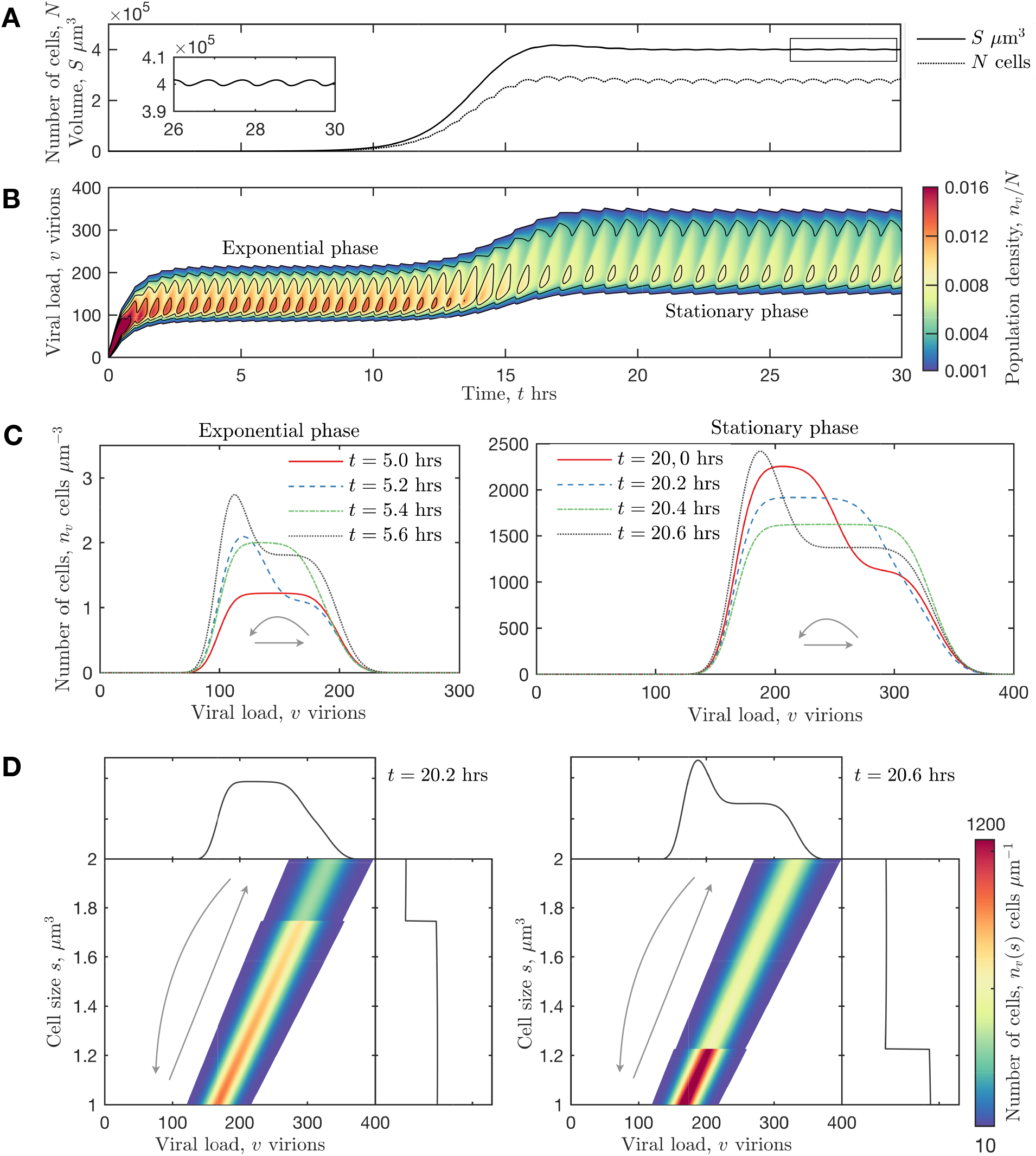
A numerical solution to equations (1) and (**??**) for the number of cells *n_v_* (*t, s*) with size *s* (*μ*m^3^) and viral load *v* (virions per cell) with carrying capacity *κ* = 10^6^ *μ*m^3^ = 10^−6^ mL) and an initial homogeneous cell size distribution (*σ*(*s*) = 1 for 1 ≤ *s* ≤ 2 *μ*m^3^). The parameter values are shown in Table 1. **A** The total number of cells *N* and the collective size of all cells *S* (*μ*m^3^) (defined in equation (2)). Inset plot depict the oscillations in *S*. **B** Colour-graded contour plot for the proportion of cells with *v* virion particles per cell (given by 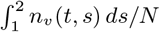). **C** Plots of the total number of cells with *v* virion particles per cell 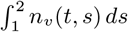 at t =5, 5.2, 5.4 and 5.6 hours (left plot) and at *t* =20, 20.2, 20.4 and 20.6 (right plot). **D** Heat map plots of the total number of *nv* (*t*, *s*) with viral load *v* particles per cell and size *s μ*m^3^ at *t* =20.2 hours (left) and 20.6 hours (right).

Figure 3B and C show the distribution of viruses among the cell population 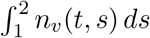 and illustrates the quasi-steady periodic state during the exponential phase and the stationary phase of population growth. During the exponential growth phase, each periodic cycle close-to doubles the number of cells with a period length of approximately 0.4 hours. From a broad uniform peak across 100 < *v* < 200, a sharp peak emerges with doubled magnitude around *v* = 100; this sharp peak slightly decays in size over time while it broadens to a uniform distribution across 100 < *v* < 200. The sharp peak is generated by the production of two daughter cells with a viral load that, on average, is half the parental viral load and the slight decay in peak magnitude is generated by cell lysis. During the stationary phase, the viral load-distribution dynamically cycles in a similar way to the exponential phase except that: (i) each cycle has a period of approximately 0.8 hours, (ii) the total number of cells after each cycle remain constant, (iii) the broad peak is distributed across 150 < *v* < 350 and (iv) the sharp peak is initially centred around *v* = 175 virions per cell. We note that the distribution of viral concentration among cells are shown in Figure **??**B and hold steady profiles during the exponential and stationary phase.

Figure 3D depicts the time-evolution of the distribution of *v* and *s* among cells *n_v_* (*t, s*) during the stationary phase of population growth. These plots illustrate how the quasi-steady state breather dynamics are produced by local increases in cell size *s* (cell growth) and viral load *v* (viral synthesis), decay in the number of cells due to cell lysis, and non-local halving (approximately) of both *s* and *v*, and cell doubling, due to cell division. The initial homogeneous distribution of cell size among cells is preserved for all time. Furthermore, the distribution of viral loads among cells holds a smoothed profile of the cell size distribution because both the cell size and viral loads are partitioned with mass conservation between cell progeny. The cell size distribution is discontinuous and the viral load distribution is smoothed because our model assumes that cell size is symmetrically halved and viral loads are random partitioned (binomial distribution) during cell division.

## 4 Analytical results and the condition for coexistence

This section presents and describes a continuum approximation of the mean-field model (nondimensionalised), which is fully developed and analysed in Appendix D. This model is analysed to find the quasi-steady state solution for the distribution *n*(*t, s, v*) of cell size *s* and viral load *v* during the exponential and stationary phase of cell population growth, and the conditions (in terms of model parameters) for cell and virus coexistence.

### 4.1 Nondimensional continuum mean-field model

The independent variables in the mean-field model (equation (1)) can be nondimensionalised by rescaling each variable as follows:

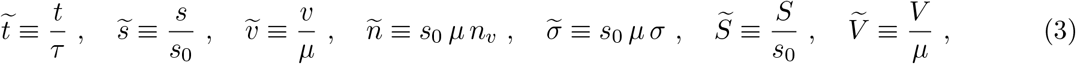

where parameter *τ* = *s*_0_/*γ* (hrs) is the mean division time (time taken for a newborn cell to divide) and *μ* (virions cell^−1^) is the mean burst size (the mean number of viruses per cell during cell lysis in the absence of cell growth and division, see Appendix B). The ranges of the independent variables are 1 ≤ *s* ≤ 2 and *v* ≥ 0. Note that the units for the number of cells remain in the model.

Burst sizes are typically large *μ* ≫ 1 (number of virions per cell). As such, by definition of the difference quotient and binomial distribution, it is reasonable to approximate virion synthesis and viral load partitioning by the following continuous functions:

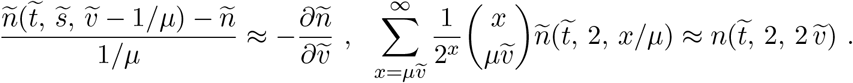

Substitution of the rescaled variables in equation (3) into equations (1) and taking the continuum limit *μ* → ∞ produces the following two-dimensional first-order hyperbolic PDE with a non-local and periodic boundary condition:

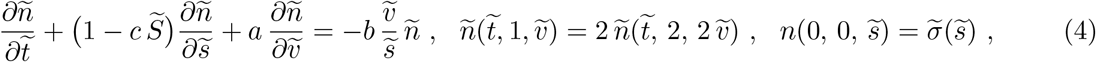

where nondimensional parameters *a*, *b* and *c* are defined as:

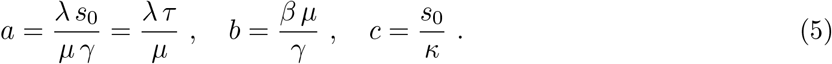

Parameter *a* is given as the product of the mean lysis time *μ*/λ (the time between initial viral synthesis and cell lysis in the absence of cell growth and division) and the mean division time *τ* = *s*_0_/*γ*. In the following subsection we show that at steady state the mean lysis time *μ*/λ (the time between initial viral synthesis and cell lysis in the absence of cell growth and division) is equal to the mean division time *τ* = *s*_0_/*γ* such that *a* = 1. Lastly, the population size *N* and the rescaled values of the collective cell volume 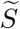 and total number of intracellular viruses 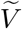 are respectively defined as:

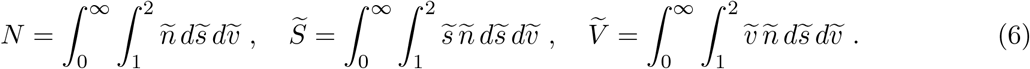

### 4.2 Quasi-steady state solution during the exponential phase of population growth

The quasi-steady state solution for the distribution of viruses and size among cells during the exponential phase of population growth can be approximated by considering an unbounded carrying capacity *κ* → ∞ (*c* = 0), which assumes no intercellular competition. From equation (4) with *c* = 0, cells that initially have the minimum size and no viral load 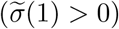 converge to the characteristic line 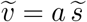 by evolving along the characteristic line 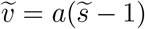 for 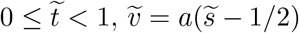 for 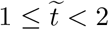, etc. The initial distribution of size among cells 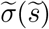is conserved for all time.

Consider the parametric solution 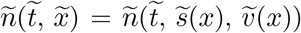 that exists only on the characteristic line:

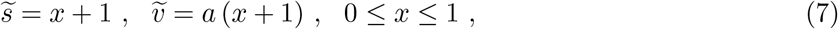

such that cells divide with a nondimensional time period of 1 (dimensional time period of *τ* hours, the minimum division time). The initial size distribution is given as 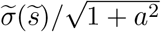 (Euclidean norm). With 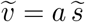, the viral concentration remains constant (at value *a*) such that 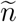 exponentially decays with rate *ab* and doubles in magnitude as it transitions from *x* = 1 to *x* = 0 (given by the periodic boundary condition stated in equation (4)):

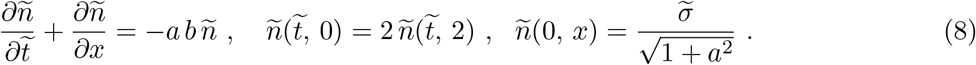

A simple application of the method of characteristics can be used to find the solution to equation (8) for 0 ≤ *t*′ ≤ 2:

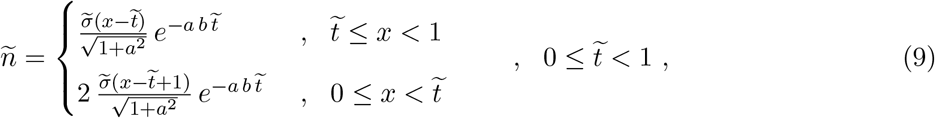

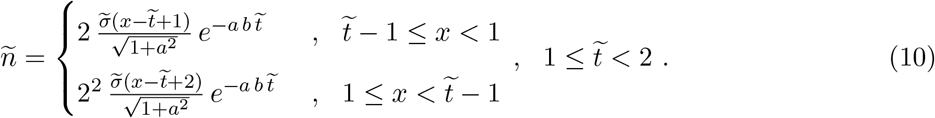

From this solution it is clear that the general solution for time 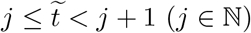:

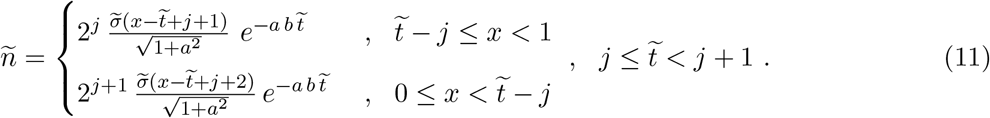

The population size *N*, volume 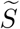 and viral content 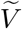 can be found by substitution of equation (11) into equation (6).

Figure 4 displays the solution to equation (8) given by equation (11) for the case where cell size and virions are uniformly distributed among cells 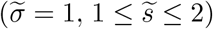. The profile of the initial distribution is conserved as the population distribution decays with rate a b, incrementally translates to large cell sizes (cell growth) and viral loads (viral synthesis) and doubles in magnitude with periodicity of 1 as it passes through the periodic boundary from *s* = 2 and *v* = 2 *a* to *s* = 1 and *v* = *a* respectively (cell division). These properties were observed in the numerical solution of the discrete mean-field model shown in Figure 3. Equation (11) and Figure 4 show that the number of cells and viruses will grow in size only if the population decay (rate *ab*) is less than population growth (periodicity of 1). As such, the condition for cell-virus coexistence is that the product of the viral synthesis rate λ (number of virions per hour), the cell lysis rate *β* (virions *μ*m^−3^ hours^−1^) and division time *τ* (hours) must be less than the product of log(2) and the cell growth rate *γ* (*μ*m^3^ hour^−1^):

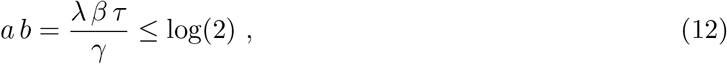

**Figure 4:**
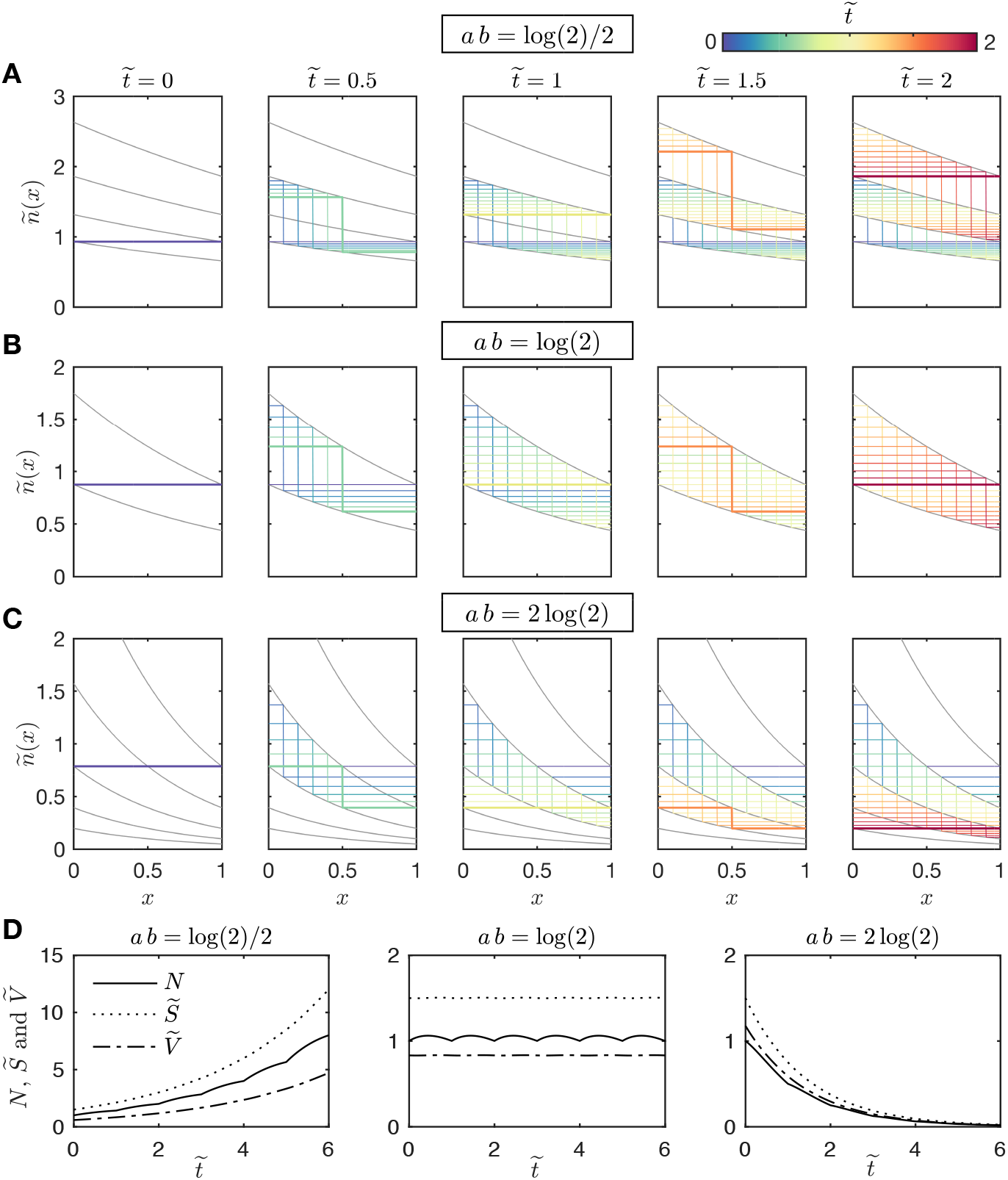
Analytical quasi-steady state solution to the nondimensionalised continuum mean field model (equation (**??**11)) for the distribution of size 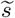 and viral loads 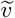 among cells 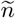) parametrised on the characteristic line *v* = *a s* (*s* = *x* +1, *v* = *a*(*x* + 1),0 ≤ *x* ≤ 1) during the exponential phase of population growth (where there is no intercellular competition *c* = 0 (*κ* → ∞)). Shown are the solutions for *a b* = log(2)/2 (**A**), log(2) (**B**) and, 2 log(2) (**C**) for times 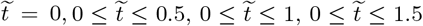 and 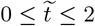 from columns left to right. The distribution at different times points are represented in different colours as indicated by the colour bar. **D** The analytical solution for the total number of cells *N* (solid line), volume 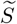 (dashed line) and total number of viruses 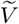 (dotted-dashed line) for parameter values *ab* = log(2)/2, log(2) and 2log(2) (from left to right).

The population size *N*, volume 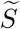 and viral content 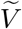 oscillate around fixed values for *ab* = log(2), grow for *ab* < log(2) and decay for *ab* > log(2). Both *N* and *S* remain at a steady value when *ab* = log(2) and the initial size distribution *σ*(*s*) is proportional to *e*^−*abs*^.

#### 4.2.1 Continuum steady state solution during the stationary phase of population growth

The steady state solution 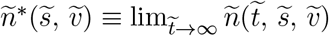 is a special case of the breather solution where the solution does not change over time during the stationary population growth phase. The equation that determines 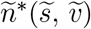 can be found by setting 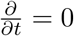 in equation (4):

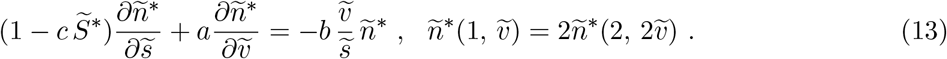

The method of characteristics can be used to find the solution 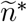 to equation (13). Let 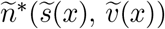 be parameterised by variable 0 ≤ *x* ≤ *T*, where *T* is the time (nondimensional) taken for a newborn cell to divide.It is clear from equation (13) that the steady state characteristic curve is given by:

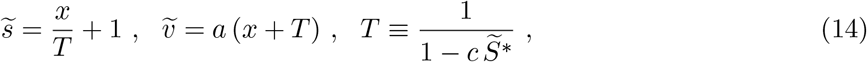

where 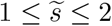 and 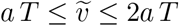. The viral concentration remains constant along the characteristic curve such that the solution to 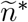 is given by an exponential function of *x*:

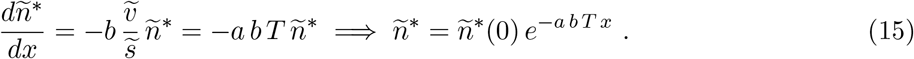

The boundary condition *n*\*(0) = 2 *n*\*(*T*) is satisfied when the division time is:

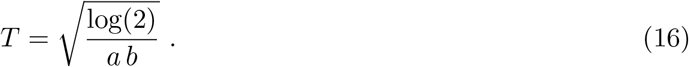

The definition of 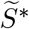 in equation (6) and (14) can be used to find the value of *n*\*(0):

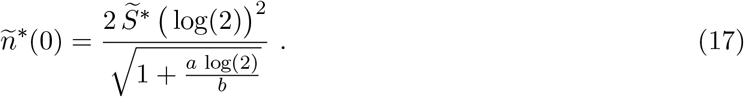

By the definition of the mean burst size (see Appendix B), equation (15) and (17) can be used to show that the value of nondimensional parameter *a* is *a* = 1:

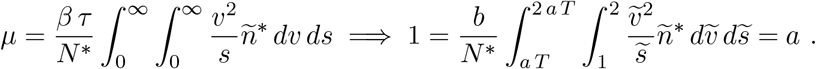

This implies that the mean burst size is given by the product of the viral production rate and the mean division time *μ* = λ*τ*. Indeed, the mean burst size has been shown to increase proportionally to the Therefore, from equation (17) the steady state solution is:

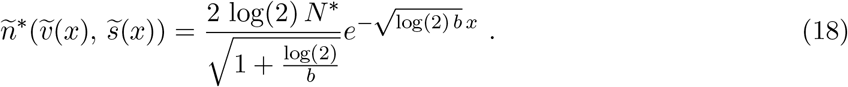

The steady state number of cells *N*\*, total population volume 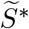 and total number of intracellular viruses 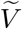 is found by substitution of equation (18) into equation (6):

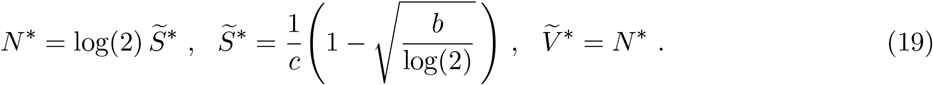

Equation(18) and (19) are plotted in Figure 5 for various values of nondimensional parameter *b*. From this analysis it is clear that the condition for cell-virus coexistence is that the product of the viral synthesis rate λ (number of virions per hour), the cell lysis rate *β* (virions *μ*m^−3^ hours^−1^) and division time *τ* (hours) must be less than the product of log(2) and the cell growth rate *γ* (*μ*m^3^ hour^−1^):

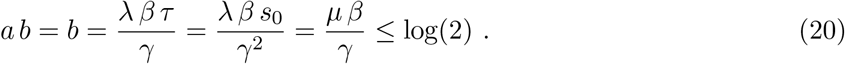

**Figure 5:**
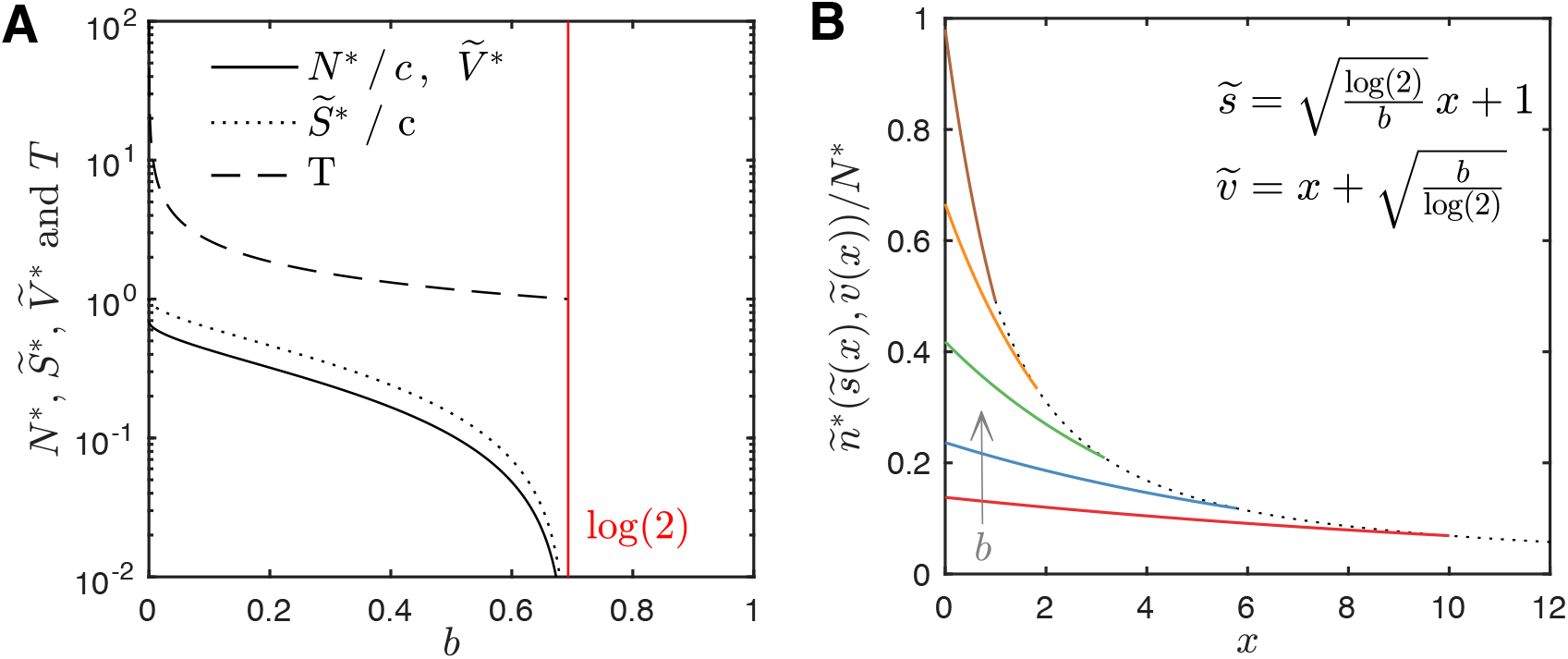
The equilibrium solution to the non-dimensionalised mean-field model (equation (4)). **A** The equilibrium population size *N*\* (solid line), viral content 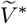 (solid line) and volume 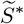 (dotted line) given by equation (19) and the division time *T* (dashed line) given by equation (16) as a function of nondimensional parameter 0 ≤ *b* ≤ log(2). **B** The parametric solution to equation (13) 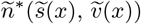, the equilibrium distribution of cell size 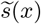 and viral load 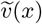 (parametric variable 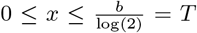) for *b* = 0.01 log(2) (red), 0.03log(2) (blue), 0.1 log(2) (green), 0.3log(2) (orange) and log(2) (brown).

Figure 5 shows that as the value of *b* changes from log(2) to 0 the population size *N*\*, the total number of intracellular viruses *V*\* (equal to the product of the mean burst size *μ* = λ *τ* (virions) and the number of cells *N*\*) and volume 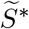 (dimensionalised by *s*_0_ *μ*m^3^) tend from 0 to *c* log(2) and *c* respectively (recall that *c* is the newborn cell size *s*_0_ *μ*m^3^ divided by the carrying capacity *κ μ*m^3^). From both Figure 5 and equation (20) we see that for fixed viral synthesis rates λ (virions hr^−1^), lysis rates *β* (virions *μ*m^−3^ hr^−1^) and newborn cell sizes *s*_0_ (*μ*m^3^), an increase in the cell growth rate γ (*μ*m^3^ hr–1) significantly promotes coexistence (cell population close to carrying capacity) by increasing both viral dilution via cell growth and viral partitioning via cell division.

We note that the viral synthesis rate λ may depend on the cell growth rate *γ* [1, 23]. If this relationship were linear (e.g. λ = *ωγ*) then the condition for coexistence is simply is that *τ* ≤ log(2)/(*βω*). This condition imposes a strict requirement for rapid cell division (small times for newborn cells to divide, *τ*), which implies that slowly dividing cells would be filtered from bacteria populations which are heterogeneous with respect to cell division rates.

## 5 Discussion

### 5.1 Summary of results

Mathematical modelling of viral-infected cells was used to study coexistence between lytic viruses and their host cells due to a coupling between the cell cycle and the lytic cycle (Figure 1). Model analysis showed that the dilution and partitioning of viral loads (the number of viruses per cell) via cell growth and division counteract viral load growth via viral synthesis, and hence cell death by lysis. As such, the cell cycle negates the lytic cycle. Generality of the proposed theory was demonstrated through the development and analysis of two different models. The model presented in the main text is representative of bacteria and assumes that cells divide once they grow (logistically) by a critical size [22] and that cell lysis rates are proportional to the viral concentration per cell. Conversely, the model presented in the Supplementary Material is representative of Eukaryotic cells and assumes that cells divide with a logistic rate following a fixed time period for cell growth and that cell lysis occurs instantaneously at a critical viral load. For both approaches, we presented numerical time-dependent solutions to equivalent stochastic and mean-field models (Figure 2 and 3), and derive analytical expressions for the distribution of viruses among cells from a continuum approximation of the mean field model (Figure 5 and 4). Model solutions were qualitatively different yet model outcomes were equivalent. Long time coexistence between lytic viruses and their host cells is set by an equilibrium between cell size growth (viral load dilution) and viral synthesis (viral load concentration), and cell division (viral load partitioning) and cell death by lysis (viral load release).

The proposed mechanism of cell-virus coexistent is conceptually similar to lysogeny (viral dormancy state). Lysogeny allows viruses to stably transfer to cell progeny as part of the host genome where it can either switch to the lytic state and kill the cell or continue to be transferred to later cell generations in the lysogenic state. Our findings suggest that viruses in the lytic state might also be transferred to cell progeny where it can either lyse cell progeny or transfer to later cell generations.

### 5.2 Ecologic effects of cell cycle-driven viral load dilution and partitioning

It is typically assumed that cells always die when infected with a lytic virus. This paradigm is supported by Lotka-Volterra models that use said assumption and predict stable cell and virus populations [1]. In these models, survival of the viral and cell population hinges on a few cells that remain uninfected during the viral spreading, and proliferate following extracellular virus decay [1]. However, predator-prey modelling often provides an incomplete and/or incorrect picture of ecological systems [27]. In this case, it is unlikely that uninfected cells remain in closed and well-mixed systems (i.e. chemostat) during viral infection cascades where cell rapidly produces tens-to-thousands of viruses which rapidly attach to cells, slowly decay [18, 28] and diffuse throughout the environment (hampered viral diffusion through structured environments such as biofilms promote cell survival [29, 30]). The present study suggests that cells and lytic viruses can coexist in the absence of uninfected cells and spatial effects due to the growth and division of infected cells, which can be observed from time-lapse videos of bacteria infected with fluorescently labelled bacteriophage [21].

Prior to the evolution of bacteriophage-resistant mutants, oscillations in the total number of cells and viruses are observed is laboratory settings (chemostats) [25]. This behaviour is characteristic of predator-prey models. In these models, the number of extracellular viruses increase as infected cells die by cell lysis and decrease to due virion decay in the presence of small numbers of uninfected cells; the number of uninfected cells proliferate when the number of extracellular viruses are small which, in turn, enables viruses to infect more cells. In our models, oscillations in the number of infected cells are due to the breather dynamics [26] of the distribution of viruses and size among cells. Breather dynamics arise from incremental increases in cell size due to cell growth and viral loads due to viral synthesis and non-local halving (approximately) of both the cell size and viral load due to cell division.

Our findings suggests that lytic viruses and their host cells might not always require evolutionary and ecological forces in order to coexist. Since cell growth and division are common properties of single cells, then their effects of viral load dilution and partitioning might be a common property of infected cells. Thus, if the dilution and partitioning of viral loads are global properties of infected cells and is sufficient for cell-virus coexistence then sufficient coexistence strategies (such as lysogeny and evolution) may not always be necessary. It is conceivable that the proposed mechanism contributes to robust and ubiquitous cell-virus coexistence by complimenting several other sufficient strategies for coexistence. For example, the coexistence of viruses and their host cells can arise through evolution by the perpetual generation of new viral-resistant cell mutants and new viral mutants that target cell mutants [8]. By establishing coexistence over short time scales, cell cycle-driven viral load dilution and partitioning might support cell and viral evolution of long time scales and relax the requirement for rapid coevolution. Furthermore, our theory states that viral infections acts as a filter for slowly dividing cells which forces rapid cell population turnover via cell division and cell death by lysis (since the infected cell population never reaches the stationary phase of population growth). As such, lytic viral infections could encourage cell evolution by forcing rapid cell population dynamics.

### 5.3 Conclusion

The present study has used mathematical modelling to suggest that the coexistence of lytic viruses and their host cells can be sufficiently maintained through perpetual viral load growth (viral synthesis), dilution (cell growth) and partitioning (cell division). Although model development was motivated by bacteria and bacteriophages, our findings might also apply to multicellular organisms. In general, our theory suggests that the cell cycle might acts as an intrinsic mechanism that enable irreplaceable, self-renewing cell populations and sub-populations (such as stem or cancerous cells) to resist population collapse in the face of a viral infection. Future work will explore the proposed theory in the human context to investigate inherent viral-resistance of self-renewing cell sub-populations such as stem cells, tumours and tissue-resident macrophages.

## 6 Appendix

### 6.1 Appendix A: Mean field model derivation via mass action kinetics

We now derive simple functional forms that model virus and cell replication via the lytic and cell cycle respectively. This derivation is complimented by the development of an equivalent stochastic model in Appendix A. Motivated by recent advances in our understanding of *E. coli* growth [22], we extend existing size-structured models of *E. coli* [20] where cell division is a function of cell size.

#### 6.1.1 Lytic cycle

In a controlled laboratory setting, *E. coli* typically synthesise mature virus particles (virions) at a constant rate following an initial delay period following infection (the eclipse phase) and up until cell death by lysis [31]. For simplicity we assume that every cell in the model is infected and produce virions at a constant rate *λ* (min^−1^):

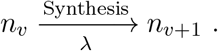

We neglect multiple infections, coupled virus synthesis and cell growth rates, and extracellular viruses. Cell death by lysis is heterogeneous with respect to viral load; the burst size is calculated from population averaging and not from a fixed threshold value of either the number or concentration of virions per cell [24]. To model cell lysis we assume that cells die at a constant rate *β* (M^−1^ min^−1^) that is proportional to the intracellular virion concentration *v*/*s* (M):

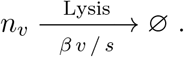

In this way, the variance in burst sizes is due to variances in both the viral loads and cell sizes.

#### 6.1.2 Cell cycle

The constant size extension theory states that bacteria divide once they grow *by* a particular size, say *s*_0_ (*μ*m^3^) [20, 22, 32–34]. Without loss of generality we assume that the minimum and maximum value of cell size can respectively be set to *s*_0_ (*μ*m^3^), the volume of newborn cells, and 2 *s*_0_ (*μ*m^3^), the volume of cells at division, such that *s*_0_ ≤ *s* ≤ 2*s*_0_. During cell division we assume that the cell size is symmetrically halved between cell progeny and that each virion is equally likely to be contained by either daughter cell (resulting in a binomial distribution) such that:

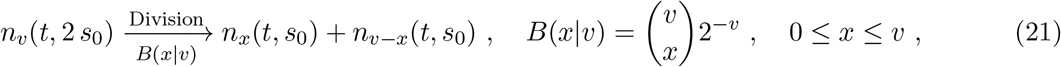

where *B*(*x*|*v*) is the binomial distribution. To incorporate logistic population growth due to cell crowding, we assume that cells grow with a maximum growth rate *γ μ*m^3^ min^−1^ that linearly decreases to zero as the total volume all cells *S* (*μ*m^3^) approaches a mamximum value *κ* (*μ*m^3^):

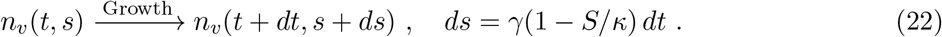

### 6.2 Appendix B: Stochastic Model derivation

Let multiset 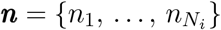 represent *N* cells with collective size *S μ*m^3^ at the discrete time point *t_i_* = *i*Δ (*i* = 0, 1,…). Each of the *N_i_* cells are given as the set *n_j_* = (*v_j_*, *s_j_ a_j_*), which quantifies the viral load 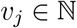 virions per cell, cell size 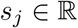 *μ*m^3^ and size grown since birth 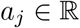 *μ*m^3^.

We now outline how we numerically simulate the stochastic model through updating 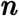 at each time step. The functional form of the stated probabilities are defined in the following section. We suppose that during time interval [*t_i_*, *t_i_* + 1] each cell has probability *P*^Synthesis^ of synthesising *k* = 0,1,… virus particles and has probability *P*^Growth^ of growing by size *x* ≥ 0 *μ*m^3^. To model cell growth and virus synthesis, at each time step we update each *n_j_* = (*v_j_*, *s_j_*, *a_j_*) by:

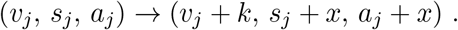

We suppose that during the time interval [*t_i_*, *t_i_* + 1], each of the cells have a probability *P*^Lysis^ of cell death by lysis and probability *P*^Division^ of cell division. These probabilities yield a set *L* cells for cell death ***n**^L^* = {*n*_*σ*_1__,…, *n_σ_L__* } and a set of *D* cells for cell division ***n**^D^* = {*n*_*μ*_1__,…, *n_μ_D__* } where 1 ≤ *σ_ℓ_*, *μ_d_* ≤ *N_i_* and *σ_ℓ_* = *σ_ℓ′_* = *μ_d_* = *μ_d_* for any ℓ, ℓ′, *d* and *d*′. To model cell death and division we update ***n*** as follows:

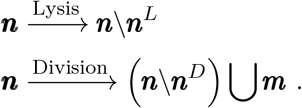

where multiset ***m*** is the set of cell progeny produced by the division of cells of set ***n**^D^*:

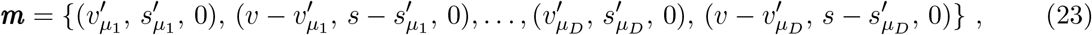

where the values 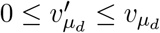 and 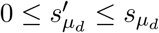 are determined by the likelihood that a parent cell with size *s* produce cell progeny with *s*′ and *s* – *s*′, *P*^*s* partition^ (*s*′|*s*), and that a parent cell with size *s* and viral load *v* produces cell progeny with *v*′ and *v* – *v*′, *P*^*v* partition^(*v*′|*v*, *s*, *s*′). These terms are derived in the following subsection.

#### 6.2.1 Probability of cell lysis

We assume that cell lysis is independent of age *a_j_* and that a cell with viral load *v_j_* has the probability of undergoing lysis during the time interval [*t*, *t* + Δ*t*] given by the following cumulative logistic function:

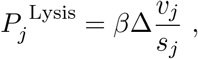

where *B* (particles per cell) is the mean burst size, *β* (per hour) is the maximum rate of cell lysis and *ϕ* (per particle per cell) is a measure of variance with respect to the burst size.

#### 6.2.2 Probability of cell division

We assume that cell division is independent of viral load *v_j_* and that a cell that has grown in size by *a_j_ μ*m^3^ has the probability of undergoing cell division during the time interval [*t*, *t* + Δ*t*] given by the following cumulative normal distribution:

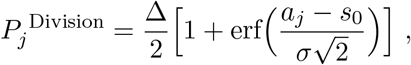

where *s*_0_ *μ*m^3^ is mean volume added between cell division events and *σ μ*m^3^ is the variance.

#### 6.2.3 Probability of synthesising *k* new virus particles

To model the synthesis of *ν* = 0, 1,… new virus particles per cell during time interval [*t*, *t* + Δ], we update each *a_j_* and *v_j_* as follows:

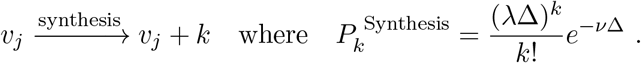

Here we assume that virus particles are created independently such that λ is determined by the Poission distribution and *γ* (per hour) is the mean synthesis rate of mature virus particles.

#### 6.2.4 Probability of growing by size *x*

Bacterial cells grow in size with a variable growth rate that is determined by the normal distribution [34]. As such, we assume that the probability that a cell grows in in size by *x μ*m^3^ during time interval Δ is given by the normal distribution:

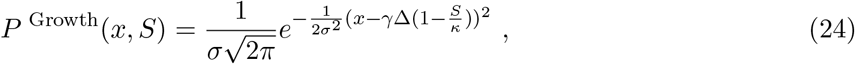

where logistic term 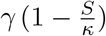 is the mean growth rate and *σ* is the growth rate variance.

#### 6.2.5 Volume and virus partitioning between daughter cells

We assume that volume is partitioned between daughter cells divide according to a normal distribution centred around half the parental cell volume:

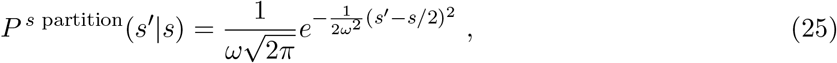

where *ω* is the division size variance. Furthermore, we assume that viral loads are partitioned between daughter cells according to the binomial distribution with probabilities equal to the daughter cell volumes:

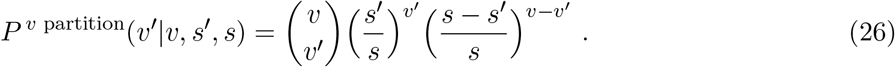

### 6.3 Appendix C: Model Parameterisation

Here we paramterise the model using typical values derived form *E. coli* and bacteriophages such as the *Escherichia virus T4* and the λ-*phage*.

#### 6.3.1 Cell growth rate *γ* and newborn cell size *s*_0_

The nutrients available to a cell governs the rate of cell growth *γ* (*μ*m^3^ hr^−1^), the newborn cell size *s*_0_ (*μ*m^3^) and hence the time between cell division *τ* (hr). Furthermore within a single cell population these values significantly vary from cell-to-cell. For *E. coli*, the population average newborn cell size and division time respectively ranging between *s*_0_ = 0.3 *μ*m^3^ and *τ* = 2 hr in nutrient poor media to *s*_0_ = 3 *μ*m^3^ and *τ* = 0.25 hr in nutrient rich media [20]. Nevertheless we assume that both the newborn cell size and the cell growth is constant such that the value of *τ* is given as:

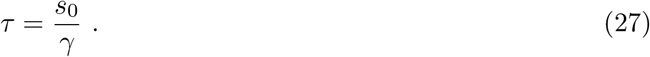

For default values we assume *E. coli* that in the absence of intercellular competition the minimum division time is *τ* = 0.5 hr and that every newborn cell has size *s*_0_ = 1 *μ*m^3^ such that the maximum growth rate is *γ* = 2 *μ*m^3^ hr^_1^.

#### 6.3.2 Viral synthesis rate λ

For *E. coli* infected with bacteriophage T4 and λ, the time period between virion production and cell lysis (the latent period) ranges from 20 to 60 mins and the mean burst size ranges from 50 to 300 virions per cell [23, 24]. Both the burst size and length of the latent period strongly depends on cell growth [1]. During this phase virion synthesis is approximately linear with respect to time. For default parameter values we consider a mean burst size of *μ* = 150 virions per cell and viral synthesis rate of λ = 200 virions per hr, which corresponds to a latent period of 45 mins.

#### 6.3.3 Cell lysis rate *β*

To estimate the cell lysis rate we calibrate our model to the burst size distribution presented in Ref. [24]. To find burst size distribution form our model we consider the time evolution of the number of viruses per cell in the absence of cell growth such that:

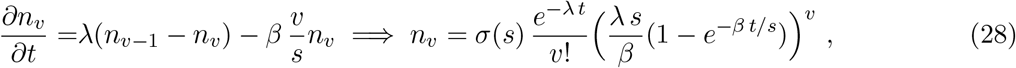

The burst size distribution *B_v_* is defined as:

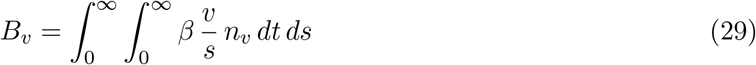

Integrating equation (28) across *t* yields:

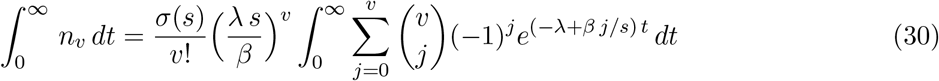

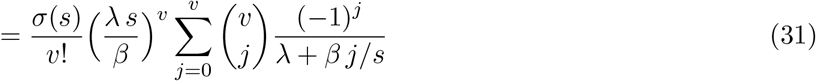

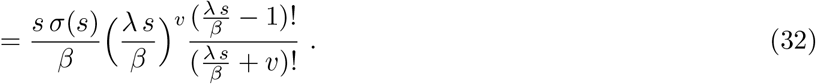

For a homogeneous size distribution where 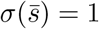 and *σ*(*s*) = 0 for 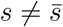 the burst size distribution is given as:

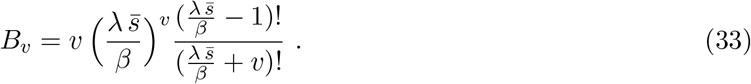

Figure 6 displays a comparison between the burst size distribution in *E. coli* measured by Delbruck (1945) [24] and the burst size distribution predicted by the model for the case that assumes a homogenous size distribution 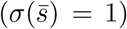 and for the case that uses a cell size distribution measured by Delbruck (1945) [24]. We find that the modelling case with a homogeneous cell size distribution sufficiently agrees with the data. Furthermore, the the mean burst size *μ* (virions per cell), defined as 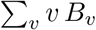 is approximately given as 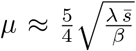. Thus, from a measured mean burst size *μ*, viral synthesis rate λ and mean size 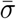, we set the lysis rate as

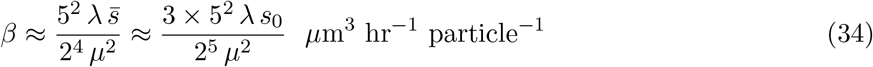

**Figure 6:**
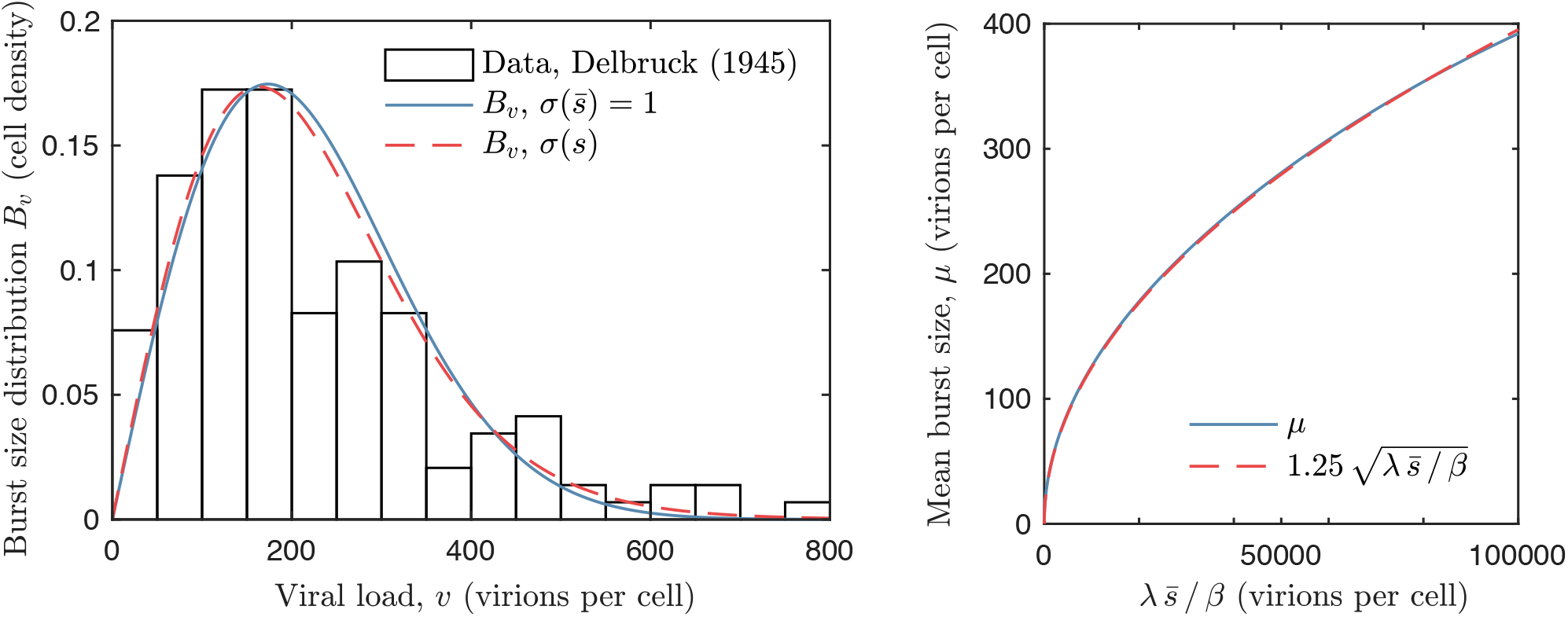
Left: the burst size distribution as presented in Ref. [24] (white bars), as predicted by equation (33) with homogeneous cell sizes 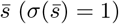 and parameter value 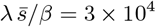 (solid blue line), and as predicted by equation (32) with cell size distribution *σ*(*s*) presented in Ref. [24] (assuming cells have a capsule shape with width 0.75 *μ*m^3^ [20]) and parameter value λ/*β* = 8.5 × 10^4^ *μ*m^−3^ (dashed red line). Right: the mean burst size 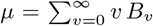 determined by equation (33) with homogeneous cell sizes for a range of parameter value 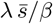 (solid blue line) and the function 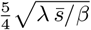 (dashed blue line).

### 6.4 Appendix D: Model analysis

We now determine the distribution of size and virions among cells during the exponential and stationary phase of cell population growth using a continuum approximation of the mean-field model.

