## Supplementary Material for "Viral-structured models of dividing cells shows cell-virus coexistence via viral load partitioning between cell progeny"

Hugh Z. Ford

#### 1 Model construction

We study the coexistence between dividing cells and intracellular lytic viruses in a well-mixed, closed and isolated system. Our construction stems from the behaviour of bacteria (e.g. *Escherichia coli*) and lytic bacteriophages (e.g. *Escherichia virus T4*, which are incapable of lysogeny). To study the relationship between the cell cycle and lytic cycle in isolation, we neglect the various evolutionary, ecological and biochemical processes that influence virus-cell coexistence. As such, our model considers a single cell and virus species and only features cell division, the partitioning of viral loads between cell progeny, subcellular virus replication and viral load-dependant cell lysis (see Figure **??**).

Our model assumptions are as follows. We assume that all cells are initially infected and neglect further infection by extracellular viruses. We assume that infected cells produce mature virus particles at a constant rate and subsequently die by lysis when the number of viruses per cell (the viral load) reaches a threshold value (the burst size). The death of subcellular viruses is neglected. We model cell division based on the concept of the constant size extension where there us a necessary cell growth period that sets a minimum time between parental and progeny division [1–4]. After this time period, we suppose that cells divide with a logistic growth rate (for a set carrying capacity) which, by mass conservation, splits the viral load between daughter cells. We now consolidate this verbal model construction with the development of an equivelant stochastic and mean field model.

##### 1.1 Statement of the stochastic model

Let $n = \{(a_1, v_1), \ldots, (a_N, v_N)\}$ represent a set of $N$ cells with age $a_j \in \mathbb{R}$ hours and viral load $v_j \in \mathbb{N}$ $(1 \leq j \leq N)$ virus particles per cell at time $t \geq 0$ hours. We assume that cell lysis is independent of age $a_j$ and that a cell with viral load $v_j$ has the probability of undergoing lysis during the time interval $[t, t + \Delta t]$ given by the following cumulative logistic function:

$$P_j^{\text{lysis}} = \frac{\beta \Delta}{1 + e^{-\phi(v_j - B)}} \ ,$$

where $B$ (particles per cell) is the mean burst size, $\beta$ (per hour) is the maximum rate of cell lysis and $\phi$ (per particle per cell) is a measure of variance with respect to the burst size. We assume that cell division is independent of viral load $v_j$ and that a cell with age $a_j$ has the probability of undergoing cell division during the time interval $[t, t+\Delta t]$ given by the following cumulative logistic function multiplied by a logistic growth term:

$$P_j^{\text{division}} = \frac{\alpha\Delta\big(1-\frac{N}{\kappa}\big)}{1+e^{-\psi(a_j-A)}} \ ,$$

where $A$ (hours) is the mean timespan of the cell growth phase $\alpha$ (per hour) is the maximum cell division rate, $\psi$ (per hour) is a measure of the variance with respect to the growth period and $\kappa$ is the cell carrying capacity.

To numerically simulate the stochastic model we choose a time step size $\Delta$ (hours) such that $P_j^{\text{lysis}} + P_j^{\text{division}} < 1$ for all $1 \le j \le N$. At each time step and for each cell we numerically generate a random number $0 < X_j < 1$. If $X_j > P_j^{\text{lysis}} + P_j^{\text{division}}$ then the cell does not divide or die via lysis. If $X_j < P_j^{\text{lysis}}$ (cell lysis) then we remove cell $j$ by setting $N \to N-1$ and:

$$n \xrightarrow{\text{lysis}} n \backslash \{(a_j,\, v_j)\} \ .$$

This operation removes the dead cell from the population. If $P_j^{\text{lysis}} < X_j < P_j^{\text{lysis}} + P_j^{\text{division}}$ (cell division) then we create two daughter cell from parental cell $j$ by setting $N \to N+1$ and:

$$n \xrightarrow{\text{division}} \Big(n \backslash \{(a_j,\, v_j\}\Big) \bigcup \{(a_j',\, v_j'),\, (a_{N+1},\, v_{N+1})\} \ ,$$

where $v_{N+1} = v_j - v_j'$ and $v_j'$ is determined from the following binomial distribution:

$$v_j' \sim \text{Bin}(v_j, 0.5) \implies P\big(v_j' = k\big) = \binom{v_j}{k}\frac{1}{2^{v_j}} \ .$$

which assumes that each virus particle is equally likely to be contained by either daughter cell. This operation removes the dividing cell from the population and replaces it with two daughter cells that share the parental viral load. To model the synthesis of $\nu = 0, 1, \ldots$ new virus particles per cell during time interval $[t, t+\Delta]$, we update each $a_j$ and $v_j$ as follows:

$$(a_j, v_j) \xrightarrow{\text{synthesis}} (a_j + \Delta, v_j + \nu) \quad \text{where} \quad \nu \sim \text{Pois}(\gamma\Delta) \implies P(\nu = k) = \frac{(\gamma\Delta)^k}{k!}e^{-\gamma\Delta} \ .$$

Here we assume that virus particles are created independently such that $\nu$ is determined by the Poission distribution and $\gamma$ (per hour) is the mean synthesis rate of mature virus particles. Numerical simulations of this model is shown in Figures 2-5 and discussed in Section 3.

#### 1.2 Statement of the mean-field model

The stochastic model can be expressed as a discrete-structured mean field model via the following infinite system of first order hyperbolic partial differential equations:

$$\frac{\partial n_v}{\partial t} + \underbrace{\frac{\partial n_v}{\partial a}}_{\text{Cell aging}} = \underbrace{\gamma(n_{v-1} - n_v)}_{\text{Virus production}} - \underbrace{\frac{\alpha\big(1-\frac{N}{\kappa}\big)n_v}{1+e^{-\psi\,(a-A)}}}_{\text{Cell division sink}} - \underbrace{\frac{\beta n_v}{1+e^{-\phi\,(v-B)}}}_{\text{Cell lysis sink}} \ , \qquad (1)$$

with boundary condition:

$$n_v(t,0) = \underbrace{2\alpha\big(1-\frac{N}{\kappa}\big)\sum_{j=v}^{\infty}\frac{1}{2^j}\binom{j}{v}\int_0^{\infty}\frac{n_j(t,a)}{1+e^{-\psi(a-A)}}\,da}_{\text{Cell division source}}\ , \tag{2}$$

where $n_v(t,a)$ is the number density of cells that contain $v = 0, 1, 2, \ldots$ virus particles at time $t \geq 0$ and $N = N(t)$ is the total number of cells. Equation (1) is closed with initial condition:

$$n_0(0,0) = N_0\ , \quad n_v(0,a) = 0 \ \text{ for } \ v,\, a > 0\ , \tag{3}$$

which represents a state where there are initially $N_0 > 0$ cells with no viral load $v = 0$ and age $a = 0$. We will either consider the initial state where $N_0 = 1$ cell or $N_0 = \kappa$ cells. The total number of cells $N$ and virus particles $V$ are defined as $N = \sum_{v=0}^{\beta} n_v$ and $V = \sum_{v=0}^{\beta} v n_v$ respectively such that their time evolution is described by:

$$\frac{dN}{dt} = \alpha\Big(1-\frac{N}{\kappa}\Big)\sum_{v=0}^{\infty}\int_0^{\infty}\frac{n_v}{1+e^{-\psi(a-A)}}\,da - \beta\sum_{v=0}^{\infty}\int_0^{\infty}\frac{n_v}{1+e^{-\phi(v-B)}}\,da\ , \quad N(0) = N_0 \tag{4}$$

$$\frac{dV}{dt} = \gamma N - \beta\sum_{v=0}^{\infty}\int_0^{\infty}\frac{v\,n_v}{1+e^{-\phi(v-B)}}\,da, \qquad V(0) = V_0\ . \tag{5}$$

Numerical solutions of equation (1)-(4) are shown in Figures 3 and 5 and discussed in Section 3.

##### 1.3 Model parameterisation

We choose default parameter values based on *E. coli* and Bacteriophage *T4*.

*E. coli*: the minimum time between cell division events is 20 to 30 minutes [5] [1–3]. Thus the observed division time is given by $A + 1/\alpha$. Thus a value $\alpha = 0$ represents an environment which cells cannot grow and $1/\alpha = 0$ represents an environment which enables cells to divide instantaneously after the growth phase.

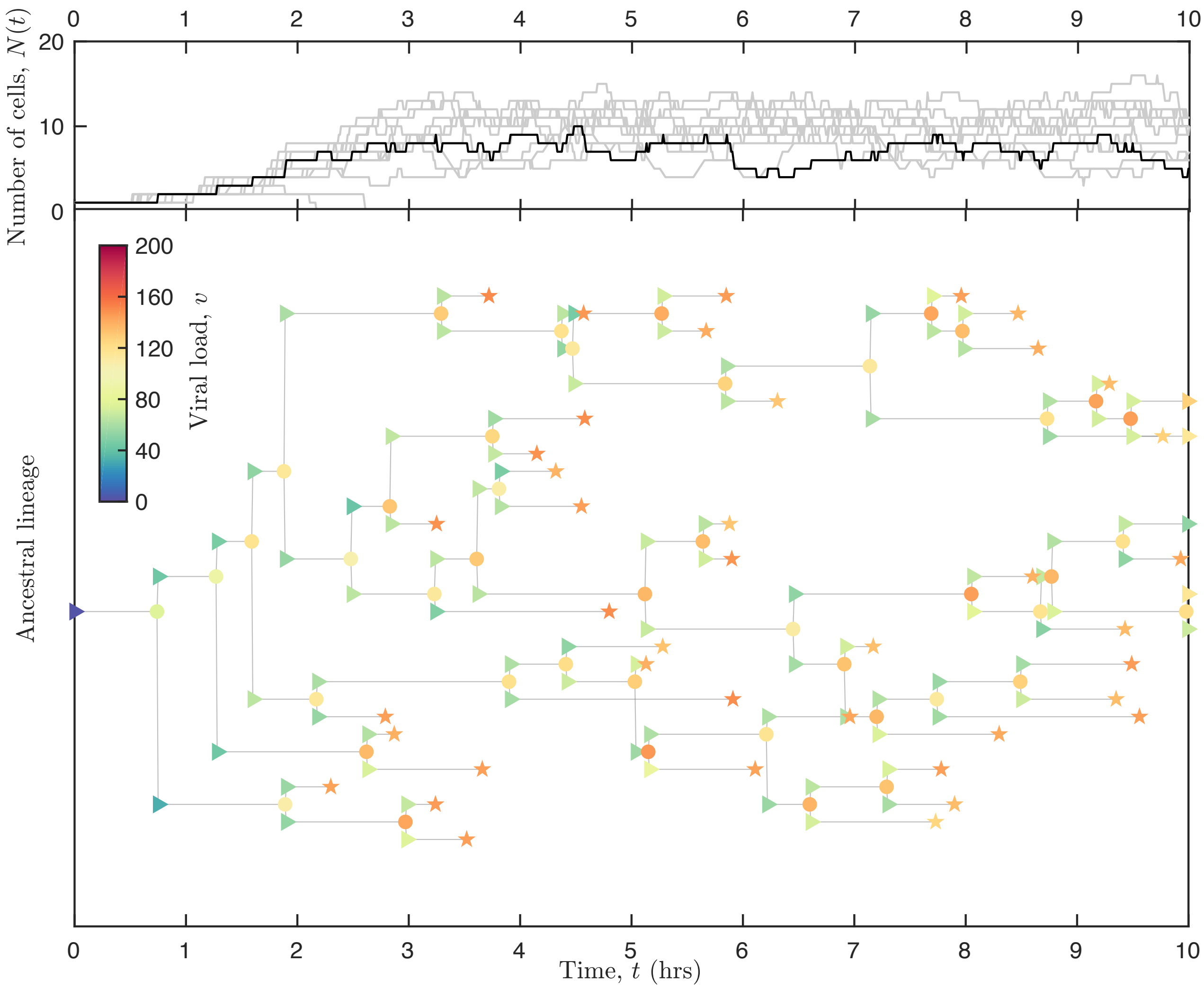


Figure 1: Simulations of the stochastic model for initially $N_0 = 1$ cell, a minimum division time of $A = 0.5$ hours, maximum division rate of $\alpha = 10$ per hour, variance in division time $\psi = 20$ per hour, carrying capacity of $\kappa = 16$ cells, burst size of $B = 150$ cells, maximum lysis rate of $\beta = 100$ per hour, mean virus synthesis rate $\gamma = 100$ particles per hour and variance in lysis time $\phi = 0.25$ per particle. Shown are the number of cells $N$ over $t = 5$ hours for 20 simulations. Also shown are the cellular ancestral lineage for scenarios where the cell population survives (labelled A) and dies (labelled B) following the infection. Each node represents the parent (circle) and daughter cells (triangle) for a division event or cell lysis (star). The colour of these nodes represent the number of viruses per cell, as indicated by the colour-bar.

#### 2 Numerical time-dependent solutions for cell-virus coexistence

We now present numerical simulations of the stochastic model and numerical solutions to the mean field model (equation (1)-(4)) using the default values shown in Table 1. These values are representative of *E. coli* infected with phage-$\lambda$. The mean field model was solved using the Crank-Nicolson discretisation method.

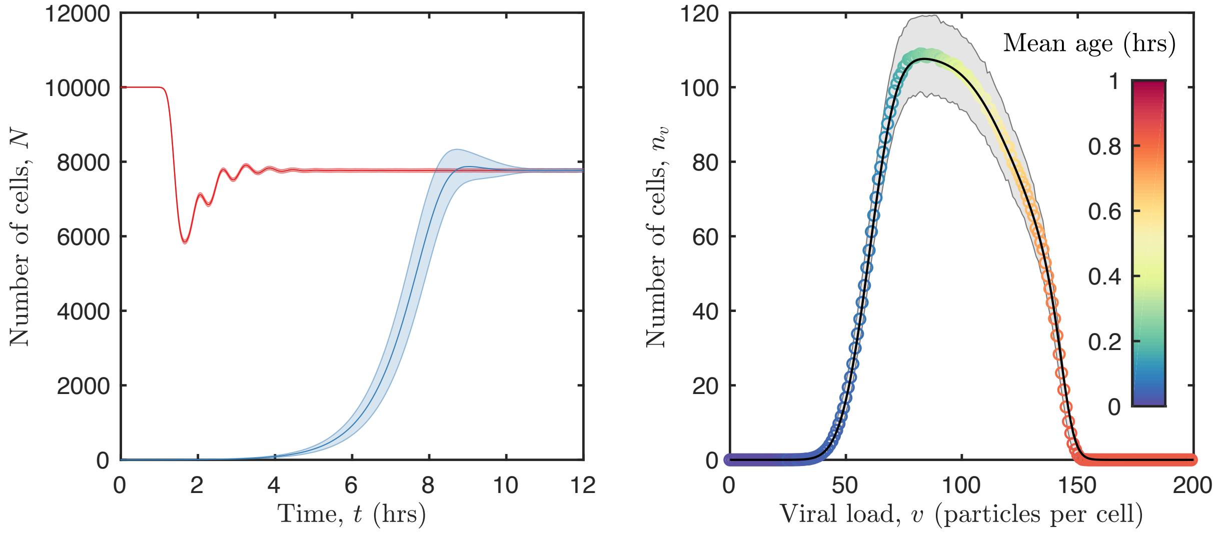


Figure 2: Numerical simulations of the stochastic model. The output shown is derived from $10^4$ runs using the same parameter values as Figure 2, except that the carrying capacity is now $\kappa = 10^4$ cells. Left: the average (solid line) and standard deviation (light area) of the total number of cells $N(t)$ when there is initially $N(0) = 1$ cell (red) and $N(0) = 10^4$ cells, the carrying capacity (blue), that contain no virus particles (i.e. $n_v(0) = 0$ for all $v > 0$) and have age $a = 0$. The numerical solution of the mean field model (equation (1) with equivalent parameter values is indistinguishable from the mean output of the numerical simulations. Right: the long time ($t = 12$ hours) total number of cells with viral load $v$ (number of particles per cell) determined from the mean field model (equation (1), black solid line) and the stochastic model (same for both initial conditions) shown as circles with a colour indicating the mean age (blue $a = 0$ to red $a = 1$ hour).

Figure 1 depicts output generated from the stochastic model for small numbers ($\kappa = 16$) of cells. The time evolution of the total number of cells $N$ is shown from 10 simulations together with a typical ancestral lineage from a single simulation. The ancestral lineage tracks the viral load $v$ of each cell (represented by node colour), the cell age (represented as the edge lengths), cells division (represented as a fork in the lineage) and cell death by lysis (represented as a terminal end). As such, the lineage illustrates how the partitioning of viral loads between cell progeny allows cell populations to persist in the face of a lytic virus infection. The random partitioning (given by a binomial distribution) of the parental viral load between daughter cells produces cell-to-cell differences which, together with stochastic viral production, can lead to different outcomes (i.e. death or division) for daughter cells.

Figure 2 shows the mean and variance in the solution from $10^3$ simulations of the stochastic model for large numbers ($\kappa = 10^4$) of cells. Figures 3 and 4 depict the numerical time-dependent solution of the mean field model. As expected, the mean output of the stochastic model was indistinguishable from the mean field model. From both the time evolution of the total number of cells $N(t)$, and the distribution of viruses and age among cells $n_v(t, a)$ at $t = 12$ hours (similar for both initial conditions),

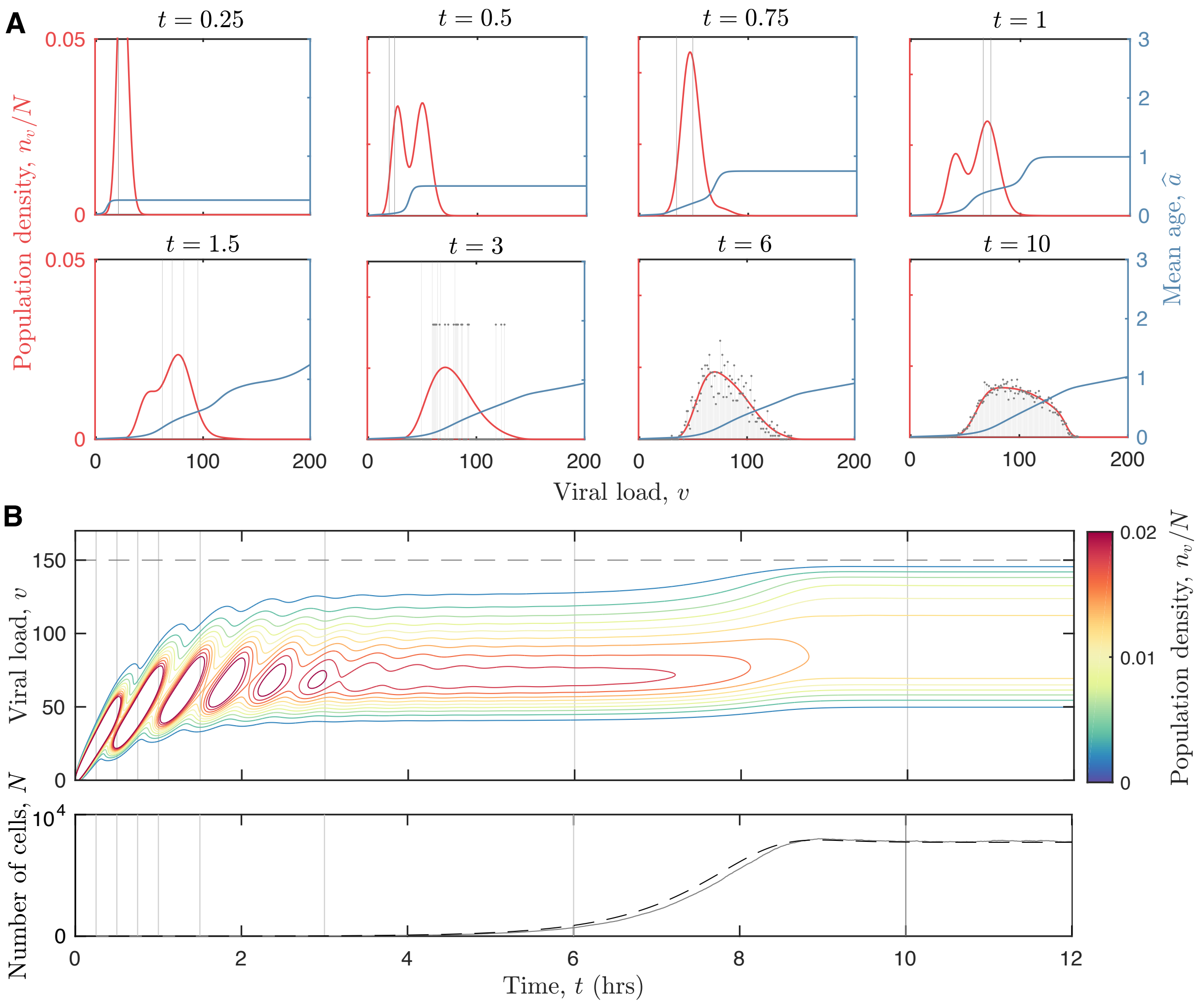


Figure 3: Numerical solution to the mean field model (equation (1)-(4)) using the same parameter values shown in Figure 2 and an initial condition of a single cells with age $a = 0$ and viral load $v = 0$. A: plots of the population density $n_v/N$ (red line) and the mean age (blue line) $\hat{a} = \int_0^t a n_v(t,a)\,da / \int_0^t n_v(t,a)\,da$ at times $t = 1,\ 1.25,\ 1.5,\ 1.75,\ 2,\ 2.25,\ 3$ and 6 hours, as indicating by the plot titles. Also shown is the population density distribution from a single realisation of the stochastic model (grey). B: contour plot of the population density (values denoted in different colours given by the colour bar) stratified by time $t$ and viral load $v$ together with the burst size $v = B = 150$ particles (dashed grey line), and a plot of the population size $N$ from the mean-field model (dashed black line) and a single realisation of the stochastic model (solid grey line). Feint solid grey lines indicate the time points that correspond to the plots in A.

it is clear that the infected cell population tends to the same equilibrium state regardless of the initial condition. Changes to the solution after 12 hours are insignificant. That is, both the stochastic and mean-field model exhibit robust cell-virus coexistence for the default parameter values for E. coli and $\lambda$-phage (Table 1).

Altogether these results show how virus-infected cell populations tend to an equilibrium state.

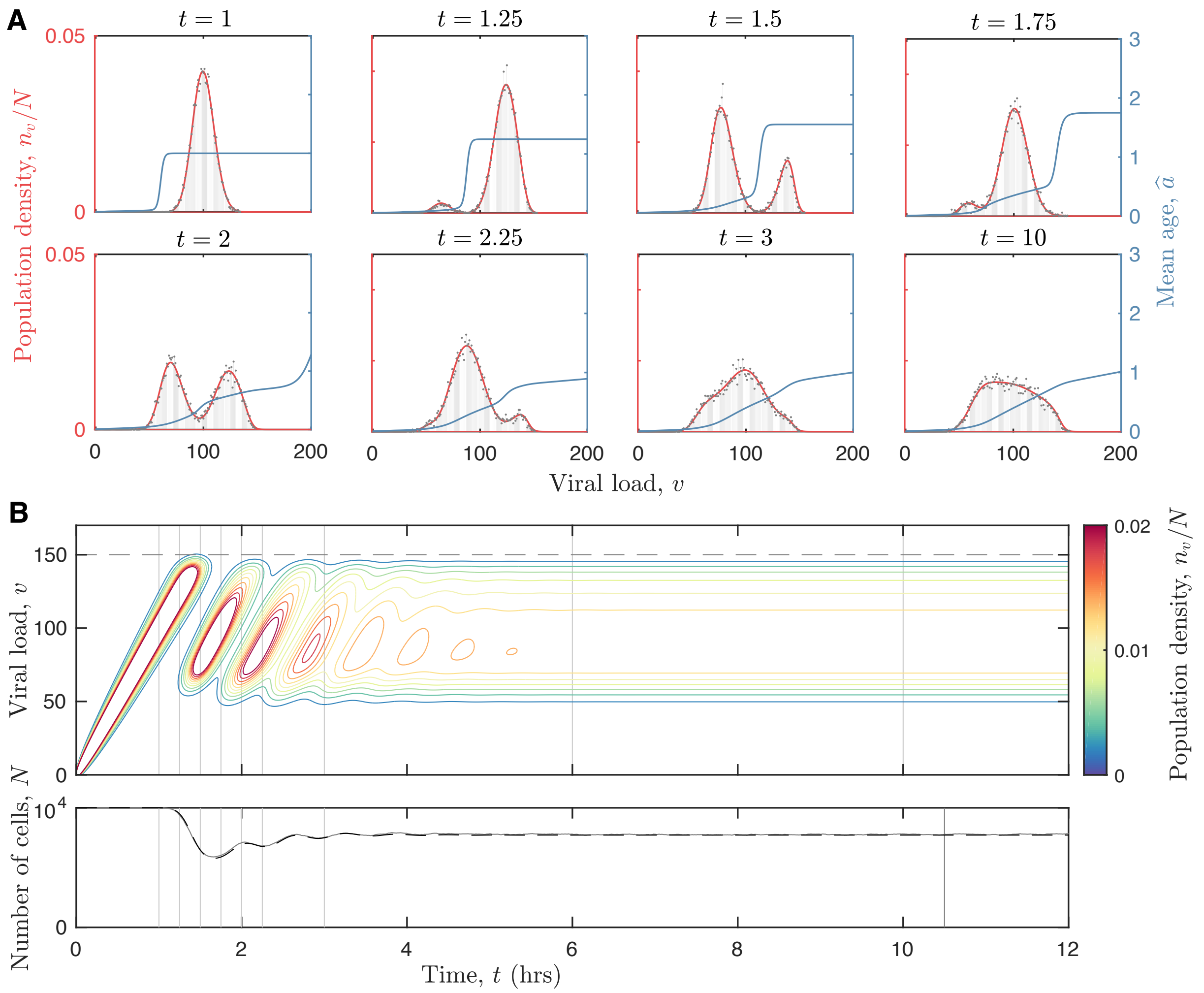


Figure 4: Same as Figure 4 except that the initial number of cells are at carrying capacity.

From the initial state, the distribution of viruses among cells locally shifts to higher viral loads (viral production). Once cells begin to divide, the viral load-distribution dynamically cycles between a single and a double peaked distribution, which are approximately centered around burst size ($v = B$) and half burst size ($v = B/2$). This profile represents the non-local production of daughter cells with a viral load that, on average, is half the parental viral load.

For the initial condition of a single infected cells (Figure 4), cells proliferate while the number of cells is small relative to the carrying capacity because cell division sufficiently dilutes viral loads below burst size such that the average lysis rate is negligible compared to the average division rate. Viruses and age are stably distributed away from the burst size (from approximately $t = 3$ to $t = 8$) while the population grows exponentially.

For the initial condition where the population is at carrying capacity (Figure 5), viral loads tend to burst size because intercellular competition (e.g. lack of resources and/or space) suppresses cell division. Once large numbers of cells die by lysis, a reduction in intercellular competition permits cell division and the partitioning of viral loads between daughter cells. This causes the segregation of viral loads into distinct bands and periodic waves where cell proliferation is dominant and where cell lysis is dominant. This is evident in the damped oscillations in the total number of cells and distinct peaks in the distribution of virus particles across the cell population..

At equilibrium (Figure 3), or close to it, viruses are heterogeneously distributed among cells with a broad peak that extend to higher viral loads form roughly half the burst size ($B = 150$ particles). The bulk of the cell population contain viral loads well below the burst size and have an age that is less than the lysis time ($\gamma/B = 2/3$ hours, the minimum division time was $A = 0.5$ hours). This distribution uniquely produces a balance between cell division, population growth and non-local viral load partition; intercellular competition, population decay and cell death/lysis; and virus synthesis and release via cell lysis.

### 3 Analysis of long time virus-cell coexistence

In this section we derive analytical expressions for the quasi-steady state distribution of viruses among dividing cells (those with $a > A$) and the steady state distribution of viruses and age among all cells. From this solution we determine the conditions for cell-virus coexistence in terms of model parameter values.

#### 3.1 Analytical model construction

The discrete-structured mean field model, equations (1) and (2), is now simplified into a continuous-structured mean field model that permits mathematical analysis.

##### 3.1.1 Model simplification

We assume that there is no variance in $v$ with respect to the burst size $B$ and no variance in $a$ with respect to the division time $A$ such that $\phi, \psi \to \infty$. We further assume that cell lysis is instantaneous when the cell viral load reaches burst size $B$ such that $\beta \to \infty$. These assumptions ensure that cells can only contain viral loads in the finite range $0 \leq v < B$. Setting $\beta$, $\phi$, $\psi \to \infty$ in equation (1) produces the following system of $B$ first order partial differential equations:

$$\frac{\partial n_v}{\partial t} + \frac{\partial n_v}{\partial a} = \begin{cases} \gamma(n_{v-1} - n_v) \ , & 0 < a < A \\ \gamma(n_{v-1} - n_v) - \alpha\big(1 - \frac{N}{\kappa}\big)n_v \ , & A \leq a \leq t \end{cases} \ , \quad 0 \leq v < B \ , \tag{6}$$

with boundary condition:

$$n_v(t, 0) = 2\alpha\big(1 - \frac{N}{\kappa}\big) \sum_{j=v}^{B} \frac{1}{2^j} \binom{j}{v} \int_A^\infty n_j(t, a)\, da \ . \tag{7}$$

Hereafter we will only consider the initial condition where there is one infected cell with age $a = 0$:

$$n_0(0, 0) = 1 \ , \quad n_v(0, a) = 0 \ , \quad v, \, a > 0 \ . \tag{8}$$

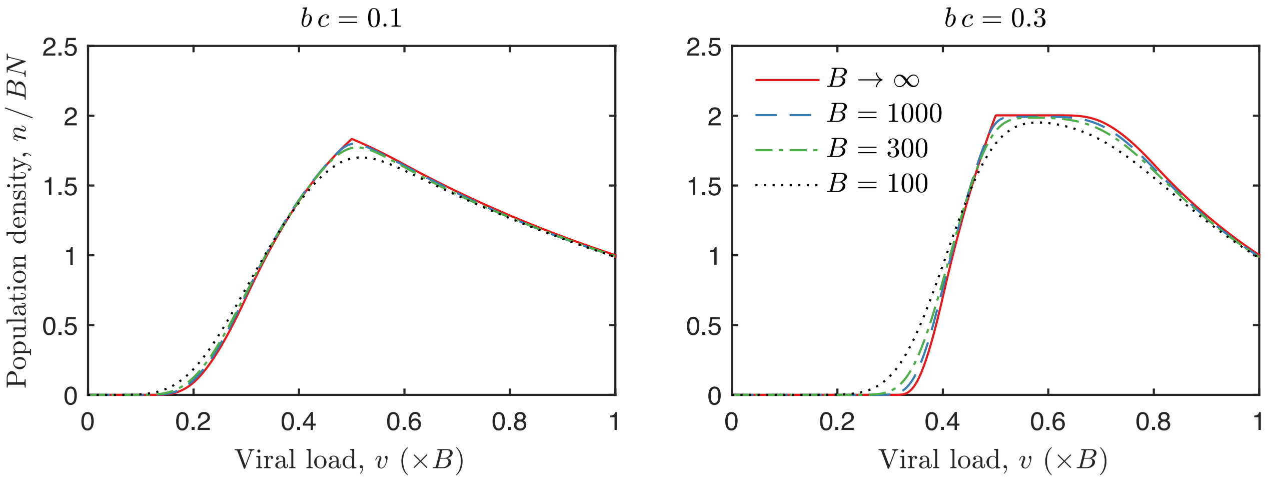


Figure 5: Numerical solutions of the the long-time density distribution of viruses among cells for burst size $B = 100$ (black dotted line), 300 (green dashed-doted line), 1000 (blue dashed line)—determined from the discrete-structured mean field model (equation (7) and (8))—and $B \to \infty$ (red solid line)—determined from the continuous mean field model (equations (9) and (10)).

###### 3.1.2 Nondimensionalisation

The simplified mean field model, equations (6) and (7), is nondimensionalised as follows:

$$\widetilde{n}(\widetilde{t},\widetilde{a},\widetilde{v}) \equiv \frac{A\,n_v}{\kappa B}\ , \quad \widetilde{N} \equiv \frac{N}{\kappa}\ , \quad \widetilde{t} \equiv \frac{t}{A}\ , \quad \widetilde{a} = \frac{a}{A}\ , \quad \widetilde{v} \equiv \frac{v}{B}\ .$$

With these new variables, equations (6) and (7) become:

$$\frac{\partial \widetilde{n}}{\partial \widetilde{t}} + \frac{\partial \widetilde{n}}{\partial \widetilde{a}} + \gamma A\Big(\widetilde{n} - \widetilde{n}(\widetilde{v} - 1/B, \widetilde{t})\Big) = \begin{cases} 0\ , & 0 \le \widetilde{a} < 1 \\ -\alpha A(1-\widetilde{N})\widetilde{n}\ , & 1 \le \widetilde{a} \le \widetilde{t} \end{cases} \tag{9}$$

$$\widetilde{n}(\widetilde{t},0,\widetilde{v}) = 2\alpha A\big(1-\widetilde{N}\big) \sum_{j=B\widetilde{v}}^{B} \frac{1}{2^j}\binom{j}{B\widetilde{v}} \int_1^{\widetilde{t}} \widetilde{n}\Big(\widetilde{t},\widetilde{a},\frac{j}{B}\Big)\, d\widetilde{a}\ . \tag{10}$$

###### 3.1.3 Continuum approximation

Burst sizes are typically large $B \gg 1$. By definition of the difference quotient and the binomial distribution:

$$\frac{\partial \widetilde{n}}{\partial \widetilde{v}} \equiv -\lim_{B\to\infty} \frac{\widetilde{n}(\widetilde{v}-\frac{1}{B},\widetilde{a},\widetilde{t}) - \widetilde{n}}{\frac{1}{B}} \quad \text{and} \quad 2\int_{\alpha A}^{\infty} \widetilde{n}(\widetilde{t},\widetilde{a},2\widetilde{v})\, d\widetilde{a} \equiv \lim_{B\to\infty} \sum_{j=B\widetilde{v}}^{B} \frac{1}{2^j}\binom{j}{B\widetilde{v}} \int_{\alpha A}^{\infty} \widetilde{n}\Big(\widetilde{t},\widetilde{a},\frac{j}{B}\Big)\, d\widetilde{a}\ ,$$

in the continuum limit $B \to \infty$ equations (9) and (10) simplify to the following three-dimensional, non-local, first order partial differential equation:

$$\frac{\partial \widetilde{n}}{\partial \widetilde{t}} + \frac{\partial \widetilde{n}}{\partial \widetilde{a}} + c\frac{\partial \widetilde{n}}{\partial \widetilde{v}} = \begin{cases} 0 & \text{for} \quad 0 < \widetilde{a} < b \\ -b\,(1-\widetilde{N})\widetilde{n} & \text{for} \quad 1 \le \widetilde{a} \le \widetilde{t} \end{cases}, \tag{11}$$

with boundary conditions:

$$\widetilde{n}(\widetilde{t},0,\widetilde{v}) = 4\,b\,(1-\widetilde{N})\int_b^\infty \widetilde{n}(\widetilde{t},\widetilde{a},2\widetilde{v})\,d\widetilde{a}\ , \quad 0 < v < \frac{1}{2} \tag{12}$$

$$\widetilde{n}(\widetilde{t},\widetilde{a},0) = 0\ , \tag{13}$$

since cells cannot obtain $\widetilde{v} = 0$ by halving alone. Here, $b$ is the product of maximum division rate $\alpha$ and minimum division time $A$ and parameter $c$ is the minimum lysis rate $\gamma/B$ multiplied by $A$:

$$b = \alpha\,A\ , \quad c = \frac{\gamma\,A}{B}. \tag{14}$$

Lastly, the time evolution of the total number of cells is given by the following ordinary differential equation:

$$\frac{d\widetilde{N}}{d\widetilde{t}} = b\,(1-\widetilde{N})\int_0^1\int_1^{\widetilde{t}} \widetilde{n}(\widetilde{t},\widetilde{a},\widetilde{v})\,d\widetilde{a}\,d\widetilde{v} - \int_0^{\widetilde{t}} \widetilde{n}(\widetilde{t},\widetilde{a},1)\,d\widetilde{a} \tag{15}$$

A comparison to the long-time distribution of viruses among cells for various viral loads $B$ is shown in Figure 5. This Figure shows how the solution to the discrete-structured mean field model converges to the solution to the continuous-structured mean field model for large values of $B$.

The tilde notation is neglected hereafter.

###### 3.1.4 Preliminary analysis of characteristic surface

A preliminary analysis of the characteristics of equations (11)-(13) is as follows. Over time $t$, the value of $a$ and $v$ for each cell change according to characteristic lines $a = t - t_0$ and $v = ca + v_0$ where $t_0$ is the time of parent cell division and $v_0$ is half the parent cell viral load (see Figure 6). Thus for an initial condition ($t = 0$) where $a = 0$ for all cells, then the solution for $n(t,a,v)$ can be written in terms of the boundary condition at $a = 0$:

$$n(t,a,v) = \begin{cases} n(t-a,0,v-c\,a) & \text{for} \quad 0 \le a \le 1\ , \quad c\,a < v < 1 \\ n(t-a,0,v-c\,a)e^{-b(1-N)(a-1)} & \text{for} \quad 1 \le a \le \frac{1}{c}\ , \quad ca < v < 1 \end{cases}\ . \tag{16}$$

Since cells divide with age $a \ge 1$, cells with viral load $v > ca + \frac{1}{2}$ will either die via cell lysis or produce progeny with $\frac{2c+1}{4} \le v < \frac{1}{2}$. Cells with viral load $0 \le v \le c$ will produce cell progeny with $\frac{c}{2} < v \le \frac{1}{2}$, these cells will produce cell progeny with $\frac{c}{4} < v \le \frac{1}{2}$, and so on. Thus the population tends to the equilibrium state $n^*(a,v)$:

$$n^*(a,v) = \begin{cases} n^*(0,v-ca) & \text{for} \ \ 0 \le a \le 1 \\ n^*(0,v-ca)e^{-b\,(1-N^*)(a-1)} & \text{for} \ \ 1 \le a \le \frac{1-c}{c} \end{cases}\ , \ c\,a + c < v \le \min(1, c\,a + \frac{1}{2})\ . \tag{17}$$

This long time solution only exists when the minimum viral load of dividing cells is less than the burst size:

$$c < \frac{1}{2} \implies \frac{\gamma A}{B} < \frac{1}{2} \tag{18}$$

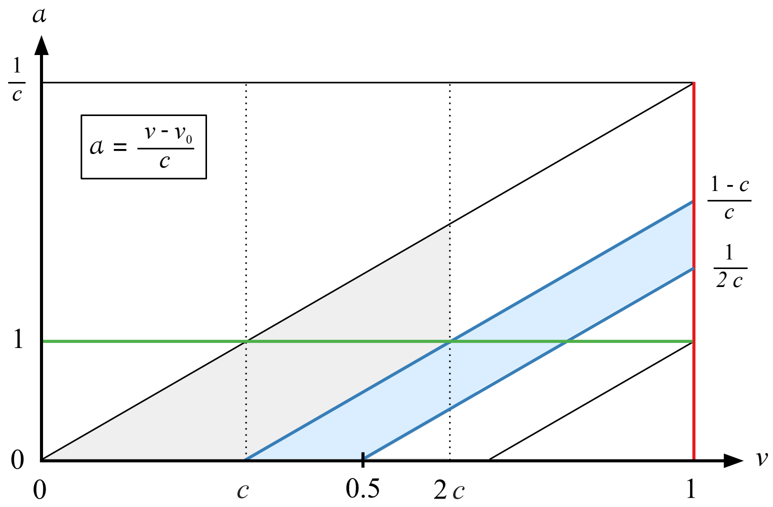


Figure 6: A schematic illustrating the characteristics of $n(t, a, v)$ across $v$ and $a$ domain: $a = (v - v_0)/c$ where $v_0$ is the viral load of cell progeny the instant after cell division. Cells divide only with $a$ and $v$ values contain within the region highlighted in red: $b \leq a \leq \frac{1}{c}$ and $ca \leq v \leq 1$. Cells at equilibrium have $a$ and $v$ values contain within the region shaded in blue: $bc \leq a \leq \frac{1}{c} - b$ and $c(a+b) \leq v \leq ca + \frac{1}{2}$. The green region indicates $a$ and $v$ values where dividing cells produce progeny that exist in the blue region.

That is, dividing cells and lytic viruses coexist only when the product of the maximum cell lysis rate $\gamma/B$ must be less half the inverse of the minimum division time $\frac{1}{2A}$.

We note that if cell division was instantaneous after the growth phase $a = A$ and unaffected by intercellular competition (i.e. $\alpha, \kappa \to \infty$), then our model predicts that cells perpetually proliferate with no cell death by lysis if equation (18) is satisfied. Guided by Figure 7, in this case we see that the population would tends to an equilibrium state where cell population evolves along the charactersic line $a = \frac{v}{c} - b$.

#### 3.2 Quasi-steady state solution for dividing cells

The quasi-steady state distribution of viruses among dividing cells (those with age $a \geq A$) is now found from the continuous-structured mean field model, equations (11)-(13).

##### 3.2.1 Time-dependent equation for dividing cells

Let $y = y(t, v)$ be the total number of cells with viral load $v > 2bc$ and age $a \geq b$:

$$y(t, v) = \int_b^{\frac{1}{c}} n(t, a, v)\, da \;, \quad 2bc < v \leq 1 \;, \tag{19}$$

such that the boundary condition for $n$, equation (12), in terms of $y$ is:

$$n(t, 0, v) = 4(1 - N)\, y(t, 2v) \;, \quad bc < v \leq \frac{1}{2} \;, \tag{20}$$

and thus, from equation (16), the time-dependent solution to $n$ for $t > \frac{1}{c}$ in terms of $y$ is:

$$n(t,a,v) = \begin{cases} 4\big(1-N(t-a)\big)y\big(t-a,0,2(v-ca)\big) & \text{for} \quad 0 \le a \le b \\ 4\big(1-N(t-a)\big)y\big(t-a,0,2(v-ca)\big)e^{-(1-N)(a-b)} & \text{for} \quad b \le a \le \frac{1}{c} \end{cases} . \tag{21}$$

Thus, from equations (11), (12), (19) and (21), the time evolution of $y$ can be expressed in closed from by the following non-local partial differential equation:

$$\frac{\partial y}{\partial t} + c\frac{\partial y}{\partial v} = 4\big(1-N(t-b)\big)\, y\big(t-b, 2(v-bc)\big) - \big(1-N\big)y \;, \quad y(t,bc) = 0 \;. \tag{22}$$

We remark that the time evolution of the total number of cells $N$, and the total number of cells with $a \ge b$, $U$, is given by the following coupled set of two ordinary differential equations:

$$\frac{dU}{dt} = 2\big(1-N(t-b)\big)U(t-b) - (1-N)U - cy(t,1) \tag{23}$$

$$\frac{dN}{dt} = \Big(1-N\Big)U(t) - cy(t,1) \;. \tag{24}$$

##### 3.2.2 Quasi-steady state assumption

Our numerical simulations shown in Figure 4 shows that soon after $t = 0$, viruses are approximately steadily distributed among cells as the population size tends to equilibrium. From this observation we separate the $y$ into two solutions:

$$y(v,t) \approx u(v)U(t) \;, \quad 2bc < v \le 1 \;, \tag{25}$$

where $u(v) = u(v; N, U)$ is the population density distribution for all cells with $a \ge b$ such that:

$$\int_{2bc}^{1} u\, dv = 1 \;. \tag{26}$$

Substituting equation (25) into equation (22) provides:

$$\frac{dU}{dt}u(v) + c\frac{du}{dv}U = 4\Big(1-N(t-b)\Big)U(t-b)\, u\big(2(v-bc)\big) - \big(1-N\big)Uu \;, \tag{27}$$

which, by substituting in equation (23), simplifies to the following delay differential equation:

$$\frac{du}{dv} = \begin{cases} -\lambda\, u(v) & \frac{1+\omega}{2} < v < 1 \\ 2\big(\lambda + u(1)\big)\, u(2v-\omega) - \lambda\, u(v) & \omega < v \le \frac{1+\omega}{2} \end{cases} \tag{28}$$

where:

$$\omega = 2\,b\,c \;, \quad \lambda = \frac{2\big(1-N(t-b)\big)\, U(t-b)}{cU(t)} - u(1) = \frac{1}{c}\Big(\frac{U'}{U} + 1 - N\Big) \;. \tag{29}$$

We note that if the number of dividing cells slowly changes relative to the number of dividing cells $U' \ll U$, then $\lambda \approx \frac{1-N}{c}$; the division rate $\alpha(1-N)$ dividing by the lysis rate $\gamma/B$. Furthermore, at equilibrium where $\frac{dN}{dt} = 0$ and $N > 0$, from equations (24) and (29) we have that $1 - N - cu(1)$ and thus $\lambda = u(1)$. As such, the value of $\lambda$ is equal to the boundary value $u(1)$ at equilibrium and, from this value, respectively increases ($\lambda > u(1)$) and decreases ($\lambda < u(1)$) when the number of dividing cells are increasing and decreasing. Furthermore, equation (24) shows that the maximum value of $U'/U$ is $U'/U = 2$ (no cell lysis or intercellular competition) and the minimum value of $U'/U$ is $U'/U = -(1-N) - cu(1)$ such that $\lambda$ is bounded by $-u(1) \leq \lambda \leq (3-N)/c$.

##### 3.2.3 Solution

It is possible to find the solution of equation (28) in terms of $u(1)$. First, for $j = 0, 1, \ldots,$ define:

$$\widehat{u}_j(v) = u(v) \quad \text{for} \quad X_{j+1} \leq v \leq X_j \ , \quad X_j = \frac{1 + (2^j - 1)\omega}{2^j} \tag{30}$$

Such that equation (28) can be expressed as the following linear system of delay differential equations

$$\frac{d\widehat{u}_0}{dv} = -\lambda \widehat{u}_0 \qquad , \quad \widehat{u}_0(1) = u(1) \tag{31}$$

$$\frac{d\widehat{u}_j}{dv} = 2\big(\lambda + u(1)\big)\, \widehat{u}_{j-1}(2v - \omega) - \lambda \widehat{u}_j \ , \quad \widehat{u}_j(X_j) = \widehat{u}_{j-1}(X_j) \ . \tag{32}$$

The solution to equation (31) is simply:

$$\widehat{u}_0 = u(1) e^{\lambda(1-v)} \ . \tag{33}$$

Starting with the solution for $\widehat{u}_0$, the solution for $\widehat{u}_j$ for all $j > 0$ can be determined recursively by using the solution to $\widehat{u}_{j-1}$ and by imposing continuity at $v = X_j = \frac{1+(2^j-1)\omega}{2^j}$. Using this method, the solution to each $\widehat{u}_j$ for $1 \leq j \leq 3$ is:

$$\widehat{u}_1 = \widehat{u}_0 + 2u(1)\Big(1 + \frac{u(1))}{\lambda}\Big)^1 \Big[ e^{\lambda\left(\frac{\omega+1}{2} - v\right)} - e^{2\lambda\left(\frac{\omega+1}{2} - v\right)} \Big]$$

$$\widehat{u}_2 = \widehat{u}_1 + 2^2 u(1)\Big(1 + \frac{u(1)}{\lambda}\Big)^2 \Bigg[ \frac{2e^{\lambda\left(\frac{3\omega+1}{4} - v\right)}}{3} - e^{2\lambda\left(\frac{3\omega+1}{4} - v\right)} + \frac{e^{4\lambda\left(\frac{3\omega+1}{4} - v\right)}}{3} \Bigg]$$

$$\widehat{u}_3 = \widehat{u}_2 + 2^3 u(1)\Big(1 + \frac{u(1)}{\lambda}\Big)^3 \Bigg[ \frac{8e^{\lambda\left(\frac{7\omega+1}{8} - v\right)}}{3 \cdot 7} - \frac{2e^{2\lambda\left(\frac{7\omega+1}{8} - v\right)}}{3} + \frac{e^{4\lambda\left(\frac{7\omega+1}{8} - v\right)}}{3} - \frac{e^{8\lambda\left(\frac{7\omega+1}{8} - v\right)}}{3 \cdot 7} \Bigg] \ .$$

A pattern emerges from which we can deduce the general formula for $\widehat{u}_j(v)$:

$$\widehat{u}_j = \widehat{u}_{j-1} + 2^j u(1)\Big(1 + \frac{u(1)}{\lambda}\Big)^j \sum_{k=0}^{j} \frac{2^{\frac{k(k-1)}{2}} e^{2^{j-k}\lambda\left(X_j - v\right)}}{(-1)^k (2;2)_k (2;2)_{j-k}} \ ,$$

where $(2;2)_k = (-1)^k \prod_{\ell=1}^{k} (2^\ell - 1)$ for $k > 0$ and $(2;2)_0 = 1$ is the q-Pochhammer symbol. Thus the solution to $u(v)$ is:

$$u(v) = u(1) \sum_{j=0}^{\infty} 2^j H(X_j - v)\Big(1 + \frac{u(1)}{\lambda}\Big)^j \sum_{k=0}^{j} \frac{2^{\frac{k(k-1)}{2}} e^{2^{j-k}\lambda\left(X_j - v\right)}}{(-1)^k (2;2)_k (2;2)_{j-k}} \ , . \tag{34}$$

where $H(x) = 1$ for $x \geq 0$ and $H(x) = 0$ for $x < 0$ is the Heaviside function. Since $u$ is a density distribution (equation (26)), integrating equation (41) across $\omega \leq v \leq 1$, and by substituting for $\lambda$ (equation (29)), provides the following algebraic equation for the boundary value $u(1)$:

$$\int_{\omega}^{1} u dv = 1 = \frac{u(1)}{\lambda} \sum_{j=0}^{\infty} \Big(1 + \frac{u(1)}{\lambda}\Big)^j \sum_{k=0}^{j} \frac{2^{\frac{k(k+1)}{2}} \Big(e^{2^{-k}\lambda(1-\omega)} - 1\Big)}{(-1)^k (2;2)_k (2;2)_{j-k}} . \tag{35}$$

We note that $2^j(X_j - \omega) = 1 - \omega$. which can be solved numerically (e.g. via Matlab's fsolve algorithm). It is of interest to understand the mean number of viruses per cell $\bar{v}$:

$$\bar{v} = \int_{\omega}^{1} v\, u(v)\, dv = \frac{u(1)}{\lambda^2} \sum_{j=0}^{\infty} 2^{-j} \Big(1 + \frac{u(1)}{\lambda}\Big)^j \sum_{k=0}^{j} \frac{2^{\frac{k(k+1)}{2}} \Big((e^{2^{-k}\lambda(1-\omega)} - 1)(2^j \lambda\omega + 2^k) - \lambda(1-\omega)\Big)}{(-1)^k (2;2)_k (2;2)_{j-k}} \tag{36}$$

##### 3.2.4 Solution description

Equations (41) and (35) (the solution to equation (28) and quasi-steady state solution to equation (22)) predict how viruses are distributed among dividing cells and are plotted in Figures 7 and 8 for various values of nondimensional parameters $\omega$ and $\lambda$ (equation (29)). Recall that $\omega$ ($\times B$ virus particles per cell) is the minimum viral load of dividing cells—those with age $a \geq b$ ($\times \alpha^{-1}$ hours)—and $\lambda$ is a measure of the population growth/decay rate. Thus, equation (41) and (35) We note that the numerical solution to equation (35) shows that the boundary value is at the equilibrium value $u(1) = \lambda$ for:

$$\lambda = \frac{1}{1-\omega} . \tag{37}$$

As such, the number of dividing cells increases while $\lambda > \frac{1}{1-\omega}$ and decreases while $\lambda < \frac{1}{1-\omega}$.

The extent of virus particle dilution via cell division can be qualitatively inferred from the profile of $u(v)$ and quantified by the proportion of cells with the viral burst size $u(1)$, shown in Figures 7 and 8. The maximum value of the population density $u(v)$ is at burst size $v = 1$ for $\lambda < 0$ and at a point that is less than or equal to the midpoint $v = \frac{1+\omega}{2}$ (minimum: $v = \omega$ and maximum: $v = 1$) for $\lambda > 0$. When the number of dividing cells are decreasing ($\lambda < \frac{1}{1-\omega}$), the profile of $u(v)$ shows that the bulk of dividing cells contain a viral load close to burst size, between $\frac{1+\omega}{2} < v < 1$. Conversely, when the number of dividing cells are increasing ($\lambda > \frac{1}{1-\omega}$), the profile of $u(v)$ shows that the bulk of the dividing cells contain a viral load close to the minimum viral load $v = \omega$, between $\omega < v < \frac{1+\omega}{2}$. This represents a state where a proliferating cell population has partitioned the virus population into viral loads that are less than burst size. In the following subsection we will consider how viruses are distributed among cells at equilibrium $u(1) = \lambda$.

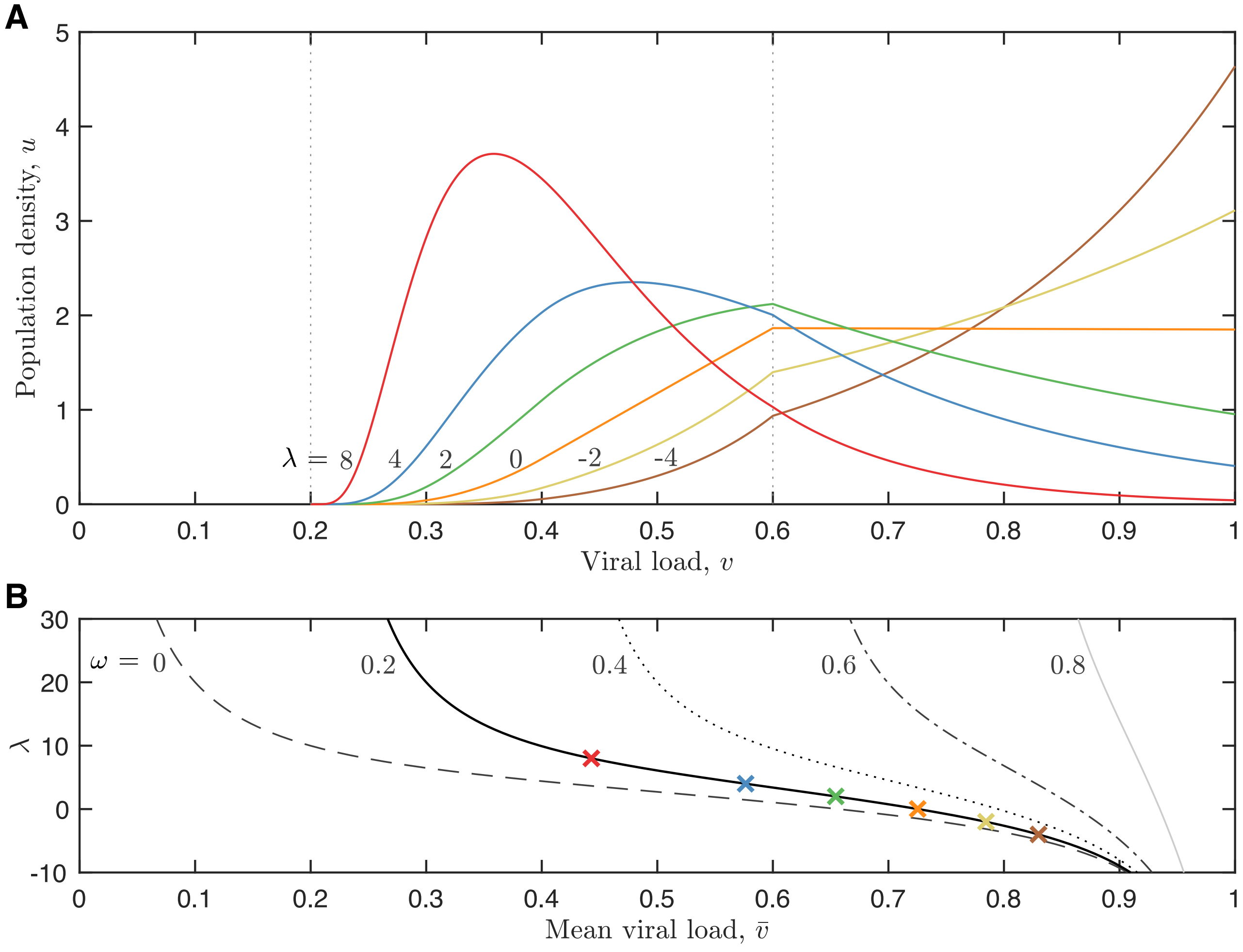


Figure 7: Plots of the quasi-steady state density distribution of viruses among dividing cells (those with age $a \geq b$), $u(v)$ (equation (41) for nondimensional parameter values $\omega = 0.2$ and $\lambda = 8$ (red), 4 (blue), 2 (green), 0 (orange), $-2$ (yellow) and $-4$ (brown), as indicated by the labelling on the right hand side of the plot. Also shown as a feint dotted lines are the values $v = \omega = 0.2$ and $v = \frac{1+\omega}{2} = 0.6$. Parameter $\omega$ is the minimum viral load of dividing cells and $\lambda$ is representative of the growth rate of dividing cells (equation (29)).

###### 3.2.5 Special case with no minimum division time $A = b = \omega = 0$

Unsurprisingly, our solution bears resemblence to the quasi-steady state solution to size-structured models of dividing cell populations (Hall and Wake, Sinko and Streifer). These models assume that cell sizes and the number of cells are unbounded:

$$y(x) = \frac{\alpha}{(2;2)_\infty} \sum_{n=0}^{\infty} \frac{2^n}{(2;2)_n} e^{-2^n \alpha x} \tag{38}$$

These models are equivalent to equation (21) for $c = 1$ and $1 - N$ To compare the solution to these models with our solution given by equations (41) and (35) we

Nondimensional parameter $\lambda$ is given by the division rate $\alpha(1 - N)$ divided by the maximum lysis

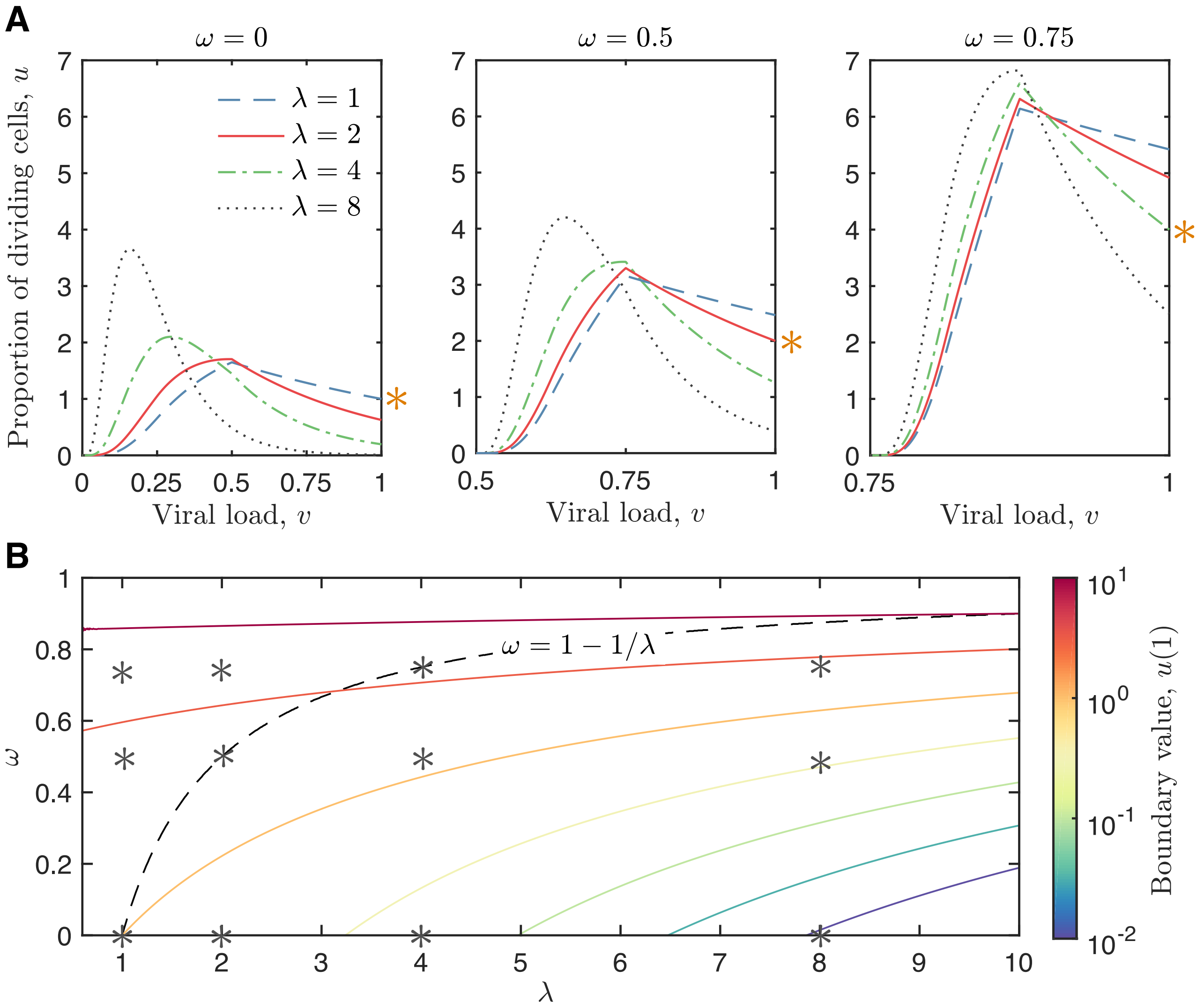


Figure 8: The distribution of viruses among dividing cells (those with age $a > A$), $u(v)$ (equation (41) for different values of nondimensional parameters $\omega$ and $\lambda$ (euaion 29). A: Plots of $u$ for $\lambda = 1$ (blue dashed line), 2 (red solid line), 4 (green dotted-dashed line) and 8 (black dotted line) and $\omega = 0$ (left), 0.25 (middle) and 0.5 (right). The curve where $\lambda = u(1)$ (the steady state value) is indicated by an orange asterisk. B: A contour plot of the boundary value $u(1)$ (equation (35)) for a range of $\lambda$ and $\omega$ values; the parameter values indicated by a grey asterisk correspond to the plots in A. The contours are $u(1) = 10^{-2}, 10^{-1.5}, \ldots, 10^1$ (blue to red). The value $\lambda = u(1)$ (the steady state value) is shown as a dashed line; an increasing number of dividing cells shifts $\lambda$ to a higher values and vice versa. The curve where $\lambda = u(1)$ is $\omega = 1 - 1/\lambda$.

rate $\gamma/B$:

$$\lambda = \frac{2(1-N)}{c} \tag{39}$$

$X_j = 2^{-j}$

$$u(v) = u(1)\sum_{j=0}^{\infty} 2^j H(2^{-j} - v)\Big(1 + \frac{u(1)}{\lambda}\Big)^j \sum_{k=0}^{j} \frac{2^{\frac{k(k-1)}{2}} e^{2^{j-k}\lambda\left(2^{-j}-v\right)}}{(-1)^k(2;2)_k(2;2)_{j-k}}\,,. \tag{40}$$

$u(1) = 0$

$$u(v) = u(1)\sum_{j=0}^{\infty} 2^j H(2^{-j} - v)\Big(1 + \frac{u(1)}{\lambda}\Big)^j \sum_{k=0}^{j} \frac{2^{\frac{k(k-1)}{2}} e^{2^{j-k}\lambda\left(2^{-j}-v\right)}}{(-1)^k(2;2)_k(2;2)_{j-k}}\,,. \tag{41}$$

##### 3.3 Equilibrium solution

###### 3.3.1 Equilibrium distribution of viruses and age among all cells

We now determine the equilibrium solution to equation (11), the distribution of viruses and age among all cells $n^*(a, v)$. This solution can be found using equation (21) and equation (41), the solution for the density distribution of viruses among dividing cells $u(v)$. At equilibrium $\lambda = u(1)$ (see subsection X). Substituting $\lambda = u(1)$ into equation (35) provides the following equation for $u(1)$:

$$1 = \sum_{j=0}^{\infty} 2^j \sum_{k=0}^{j} \frac{2^{\frac{k(k+1)}{2}} \left(e^{2^{-k}u(1)(1-2bc)} - 1\right)}{(-1)^k(2;2)_k(2;2)_{j-k}}\,. \tag{42}$$

From the numerical solution to equation (42) [analytically?] we deduce that:

$$u(1) = \frac{1}{1 - 2bc}\,, \tag{43}$$

such that, from equation (24), the equilibrium number of cells $N^*$ is:

$$N^* = 1 - cu(1) = 1 - \frac{c}{1 - 2bc} = \frac{1 - c(2b + 1)}{1 - 2bc}\,. \tag{44}$$

The total number of dividing cells is $U^* = N^*(1 - 2bc)$. Thus, from equation (21), the steady state solution for $n$ in terms of $u$ (equation (42)) is:

$$n^*(a, v) = \begin{cases} 4\,c\,\frac{1-c(2b+1)}{1-2bc}\,u(2v - 2ca) & \text{if } \ 0 \le a \le b\,, \quad ca + bc < v < ca + \frac{1}{2} \\ 4\,c\,\frac{1-c(2b+1)}{1-2bc}\,u(2v - 2ca)\,e^{-\frac{c(a-b}{1-2bc)}} & \text{if } \ b \le a \le \frac{1}{c} - b\,, \quad ca + bc < v < \min(ca + \frac{1}{2}, 1) \end{cases} \tag{45}$$

where, from equation (41), $u$ is given as:

$$u(v) = \frac{1}{1 - 2bc}\sum_{j=0}^{\infty} 4^j\, H\Big(\frac{2(2^j - 1)bc + 1}{2^j} - v\Big) \sum_{k=0}^{j} \frac{2^{\frac{k(k-1)}{2}} e^{\frac{2^{j-k}}{1-2bc}\left(\frac{2(2^j-1)bc+1}{2^j} - v\right)}}{(-1)^k(2;2)_k(2;2)_{j-k}}\,,. \tag{46}$$

##### 3.4 Conditions for coexistence

Existence implies that $N^* > 0$. Thus the lysis rate must be less than the maximum division rate divided by the minumum:

$$\frac{\gamma}{B} < \frac{\alpha}{1 + 2\alpha A}\,. \tag{47}$$

If cell division can be instantaneous following the minimum division time $A$ (where $\alpha \to \infty$), then this equation becomes equivalent to equation (17).

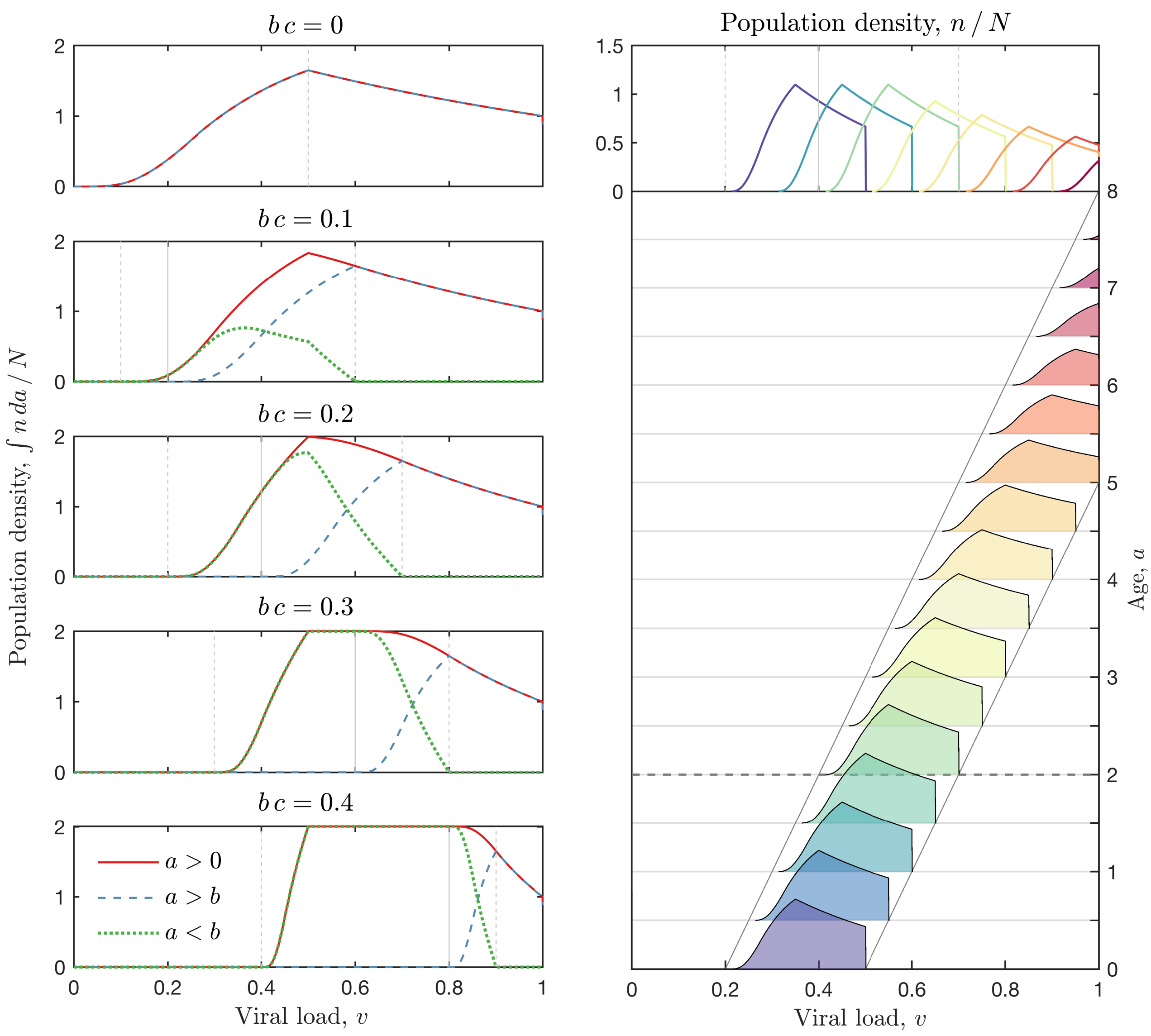


Figure 9: Analytical solution for the equilibrium distribution of viruses among cells together with the cell age $a$ (time-since-division), $n^*(a, v)$ (equation (45)). On the left set of plots, the distribution of viruses among all cells $\int_0^{\frac{1}{c}-A} n^*\,da$ (red solid line), dividing cells $\int_A^{\frac{1}{c}-A} n^*\,da$ (dashed blue line) and non-dividing cells $\int_0^{A} n^*\,da$ (dotted green line) are shown for parameter value $cA$ (the product of the lysis rate minimum division time) equal to 0, 0.1, 0.2, 0.3 and 0.4 (from top to bottom). Also shown in feint grey are the minimum and maximum viral load of all cells $v = cA$ and $v = cA + \frac{1}{2}$, and the minimum viral load of dividing cells $v = 2cA$ (the maximum is $v = 1$, the burst size). The right plot shows the distribution of viruses among cells of various ages.

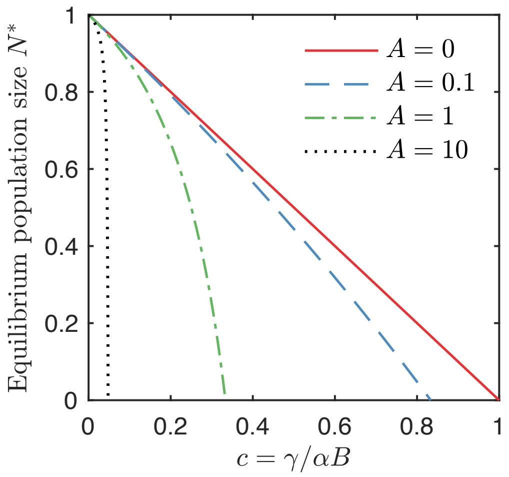


Figure 10: The equilibrium population size $N^*$


3. Campos, M. *et al.* A constant size extension drives bacterial cell size homeostasis. *Cell* **159,** 1433–1446 (2014).

4. Taheri-Araghi, S. *et al.* Cell-size control and homeostasis in bacteria. *Current biology* **25,** 385–391 (2015).

5. Cooper, S. & Helmstetter, C. E. Chromosome replication and the division cycle of Escherichia coli Br. *Journal of molecular biology* **31,** 519–540 (1968).
